# Stepping dynamics of dynein characterized by MINFLUX

**DOI:** 10.1101/2024.07.16.603667

**Authors:** Joseph Slivka, Emma S. Gleave-Hanford, Mert Golcuk, Devinda P. Wijewardena, John T. Canty, Paul R. Selvin, Mert Gur, Andrew P. Carter, Ahmet Yildiz

## Abstract

Cytoplasmic dynein is the principal motor for minus-end-directed motility and force generation functions along microtubules (MTs)^1^. Dynein converts the chemical energy of ATP hydrolysis into coordinated structural changes to step processively along MTs, but how dynein couples ATP hydrolysis to a minus-end-directed step remains controversial^2, 3^. The dynamics of dynein stepping have previously been characterized by tracking flexible regions of the motor with limited resolution^4–6^. Here, we site-specifically labeled yeast dynein at its MT-binding domain by developing a cysteine-light mutant and tracked its stepping at sub-millisecond and nanometer resolution at physiological ATP concentrations using MINFLUX^7^. We show that dynein hydrolyzes one ATP per step and takes multiples of 8 nm steps. Steps are preceded by a transient movement towards the plus end. These backward “dips” correspond to MT release upon ATP binding and subsequent diffusion of the stepping monomer around its MT-bound partner. Functional assays showed that dips terminate with a minus-end-directed movement upon ATP hydrolysis. These results provide critical insights into the order of mechanochemical events that result in a productive step of dynein.

---

Cytoplasmic dynein (dynein hereafter) is a dimeric motor that conducts minus-end-directed transport of organelles, vesicles, signaling complexes, and mRNA along MTs, as well as driving nuclear migration in developing neurons^1^. Dynein also plays central roles in mitosis, such as focusing MTs into spindle poles and pulling on astral MTs for positioning the spindle^8^. Complete knockouts of dynein stop the MT transport machinery and inhibit mitosis. Mutations that alter dynein or dynein-associated factors lead to the pathogenesis of developmental and neurological diseases^8^.

Dynein is built around a pair of heavy chains that comprise the tail and motor domains^9^ (Extended Data Fig. 1a). Each motor domain (head) is a ring of six nonidentical AAA sites (AAA1-6)^10^ that connects to an MT binding domain (MTBD) via an antiparallel coiled-coil stalk. The two rings are connected to a common tail via the linker domains. Dynein motility is primarily driven by coupling the ATPase activity at AAA1^11^ to conformational changes of its stalk/MTBD and linker^12, 13^. In addition to AAA1, AAA3 hydrolyzes ATP, and mutations to this site substantially slow dynein motility^14^. It remains uncertain whether AAA3 acts catalytically in concert with AAA1 or whether ATP hydrolysis at AAA1 alone drives each dynein step^15–17^.

ATP binding to AAA1 triggers the release of a head from the MT by altering the registry of the stalk coiled-coils^12, 18^. To generate a productive step, dynein must undergo a conformational change that biases its step size towards the minus end before the stepping head rebinds the MT. Structural studies revealed that the linker of the stepping head bends at or undocks from the surface of the AAA+ ring in the post-hydrolysis (ADP.Pi) state^19^. This “priming stroke” of the linker was proposed to move the MTBD towards the minus end^2, 20^. A recent cryo-electron microscopy study^3^ proposed an alternative model, where the linker straightens before MT binding, and a net bias in the step size is generated by swinging of the linker from AAA4 to AAA5 after the head binds the MT. Therefore, the conformational change that produces a net step in the dynein mechanochemical cycle remains controversial.

The order of events during stepping could not be directly determined from the structures of dynein captured in distinct nucleotide states^2, 3, 21^, and is better addressed by tracking the motor as it processively walks along the MT. Stepping of dynein heads has been observed by labeling the AAA+ ring with a fluorophore and tracking its position using fluorescence imaging with one-nanometer accuracy (FIONA)^4, 5, 22, 23^. Dynein moves by uncoordinated stepping of its heads, and it frequently takes steps in sideways and backward directions on the MT^4, 5^. Due to the flexibility of dynein’s structure, these measurements were affected by the thermal fluctuations and different orientations of the AAA+ ring within the dimer^6, 20, 24, 25^. The movement of dynein from one tubulin binding site to the next could only be observed at limited spatiotemporal resolution^6^. Steps also appear as instantaneous displacements at 10-100 ms temporal resolution^4, 5, 22^, and how dynein releases from the MT, moves forward, and reattaches to the MT during stepping could not be directly observed.

To determine how dynein steps along the MT lattice, we site-specifically labeled the MTBD of dynein^23^ (Fig. 1a). The artificial dimer of yeast dynein motor domain is an ideal model system to study the intrinsic stepping of dynein since it walks processively in the absence of dynein accessory proteins and has similar stepping properties to the mammalian dynein complex^17, 26^. Based on the available structures^27, 28^, 5 out of 39 cysteines are more than 5 Å exposed to the solvent (Extended Data Table 1). These cysteines were mutated to serine (Dyn_CLM_), and a single solvent-accessible cysteine was introduced at multiple candidate sites at or near MTBD for fluorescence labeling (Extended Data Table 2). We identified a mutant (Dyn_CLM_-Q3231C) that was efficiently labeled compared to Dyn_CLM_ in vitro (Fig. 1a-b, Extended Data Fig. 1b). We constructed a heterodimer of Dyn_CLM_-Q3231C (Fig. 1a, Extended Data Fig. 1c) and confirmed that this motor moves with similar velocity and stepping properties to wild-type (WT) dynein^23^ (Fig. 1c, Extended Data Fig. 2).

**Figure 1.**
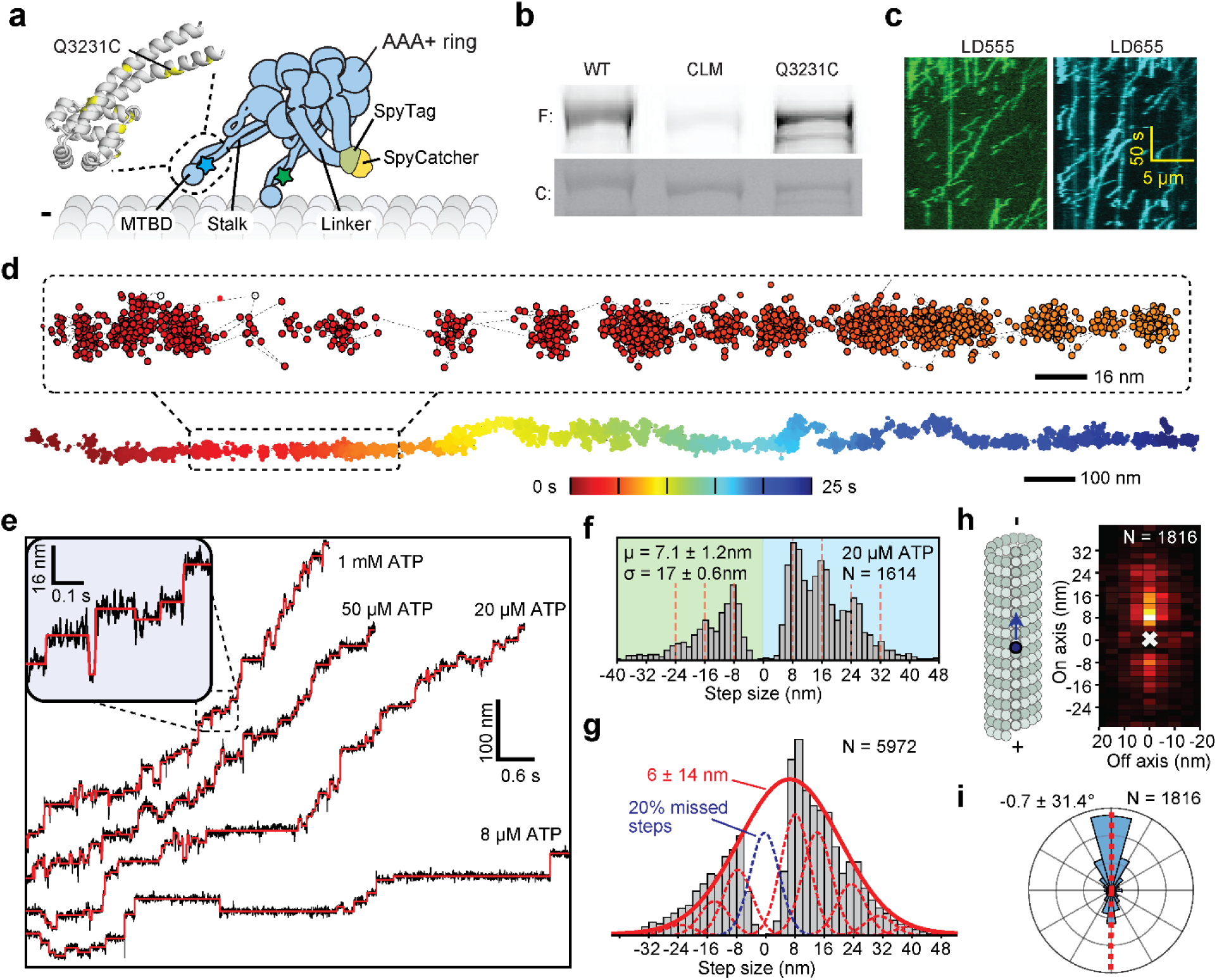
Site-specific labeling and tracking the MTBD of dynein with MINFLUX. **a,** The schematic of a heterodimer of dynein motor domains bound to the MT. (Insert) The atomic model of the MTBD shows the positions of unique cysteines (yellow) introduced after removing surface-exposed cysteines in Dyn_CLM_. Dyn_CLM_-Q3231C monomers were labeled with different dyes (blue and red stars) near their MTBD and heterodimerized via SpyCatcher-SpyTag at the tail. **b,** The denaturing gel images show that Dyn_CLM_-Q3231C is labeled with maleimide-reactive dyes more efficiently than Dyn_CLM_ (F: fluorescence, C: Coomassie-stained). **c,** Kymographs show total internal reflection fluorescence (TIRF) imaging of dynein heterodimers labeled with LD555 and LD655 dyes. **d,** An example MINFLUX trajectory of dynein in the *xy* plane, color-coded by time. **e,** Example MINFLUX trajectories of dynein stepping along the MT long axis under limiting and saturating ATP concentrations with 3.0 nm and 2.5 ms resolution. Red lines show fit to steps. **f,** Step size histogram along the MT long axis at 20 µM ATP. Red dashed lines show increments of 8 nm steps ranging from -24 nm to 32 nm. **g,** The combined step size histogram of dynein measured in 5 µM – 1 mM ATP reveals a large gap from -4 nm to 4 nm. The red curve represents a fit to a Gaussian to calculate mean ± s.d. Red dashed curves represent subpopulations fit by a Gaussian mixture model, and the blue dashed curve represents 20% “0 nm” missed steps. **h,** The short-axis step-size histogram of dynein at 20 µM ATP. **i,** The polar histogram of dynein’s stepping angle reveals no directional bias to step left or right (mean ± s.d.).

We next tracked dynein stepping at physiological (1 mM) ATP using minimal fluorescence photon flux (MINFLUX)^29, 30^ microscopy, which requires ∼100 times fewer photons than FIONA and tracks motors labeled with small (∼1 nm in size) organic dyes with sub-millisecond temporal resolution^22^. We first confirmed that MINFLUX tracking of dynein labeled at its tail exhibits stepping properties comparable to FIONA tracking at lower ATP concentrations (Extended Data Fig. 3)^4, 23, 31^. Trajectories of tail-labeled dynein exhibited large fluctuations in dye position, highlighting the importance of labeling the MT binding site to eliminate the flexibility and conformational heterogeneity of the motor. We next tracked MTBD-labeled dynein at both limiting and saturating ATP concentrations at 2.5 ms resolution (Fig. 1d-e). We observed that dynein takes increments of 8 nm steps with ∼28% of the steps taken backward (Fig. 1f). The periodicity of the dynein step size matches the distance between adjacent tubulin binding sites (8.2 – 8.4 nm)^6^. Fitting the step size histogram to a Gaussian distribution suggests a 6 nm bias to step towards the minus-end with a ±14 nm (s.d.) diffusional component to search for a tubulin binding site (Fig. 1g).

Although we cannot detect a step if the head lifts off and rebinds to the same tubulin, the step size distribution suggests that 20% of steps are “0 nm” in size and correspond to “foot stomping” of dynein (Fig. 1g; Extended Data Fig. 4). The head also frequently (27%) landed on adjacent protofilaments with nearly equal probability to step left or right^32, 33^ and these displacements on the perpendicular axis were restricted to the 25 nm diameter of MTs (Fig. 1h-i, Extended Data Fig. 5). The direction and size distribution of dynein steps were unaffected by the ATP concentration (Fig. 1f, Extended Data Fig. 4).

To determine the number of ATPs hydrolyzed by the stepping head (Fig. 2a), we analyzed the kinetics of dwell times between successive steps. Solution kinetics studies showed that the ATPase cycle of dynein is rate-limited by ADP release at saturating ATP concentrations^34^ and the motor is limited by ATP binding at low ATP. Therefore, if dynein hydrolyzes one ATP per step, stepping would be limited by a single rate constant at both limiting and saturating ATP. By contrast, it would be limited by two equal rate constants at the Michaelis-Menten constant (K_M_) of dynein for ATP (23 µM, Fig. 2b)^35^, in which ATP binding and ADP release rates become equal. In comparison, if AAA1 and AAA3 sequentially hydrolyze ATP to generate each step (Fig. 2a), the stepping kinetics can be limited by two rate constants at both limiting and saturating ATP. We found that the dwell time distribution of dynein fits best to a model with a single rate-limiting constant under both saturating (1 mM) and limiting (5-8 µM) ATP and with the convolution of two equal rate constants at 20 µM ATP (Fig. 2c, Extended Data Fig. 4d). These results show that dynein hydrolyzes one ATP per step, consistent with MINFLUX tracking of dynein stepping in neurons^17^.

**Figure 2.**
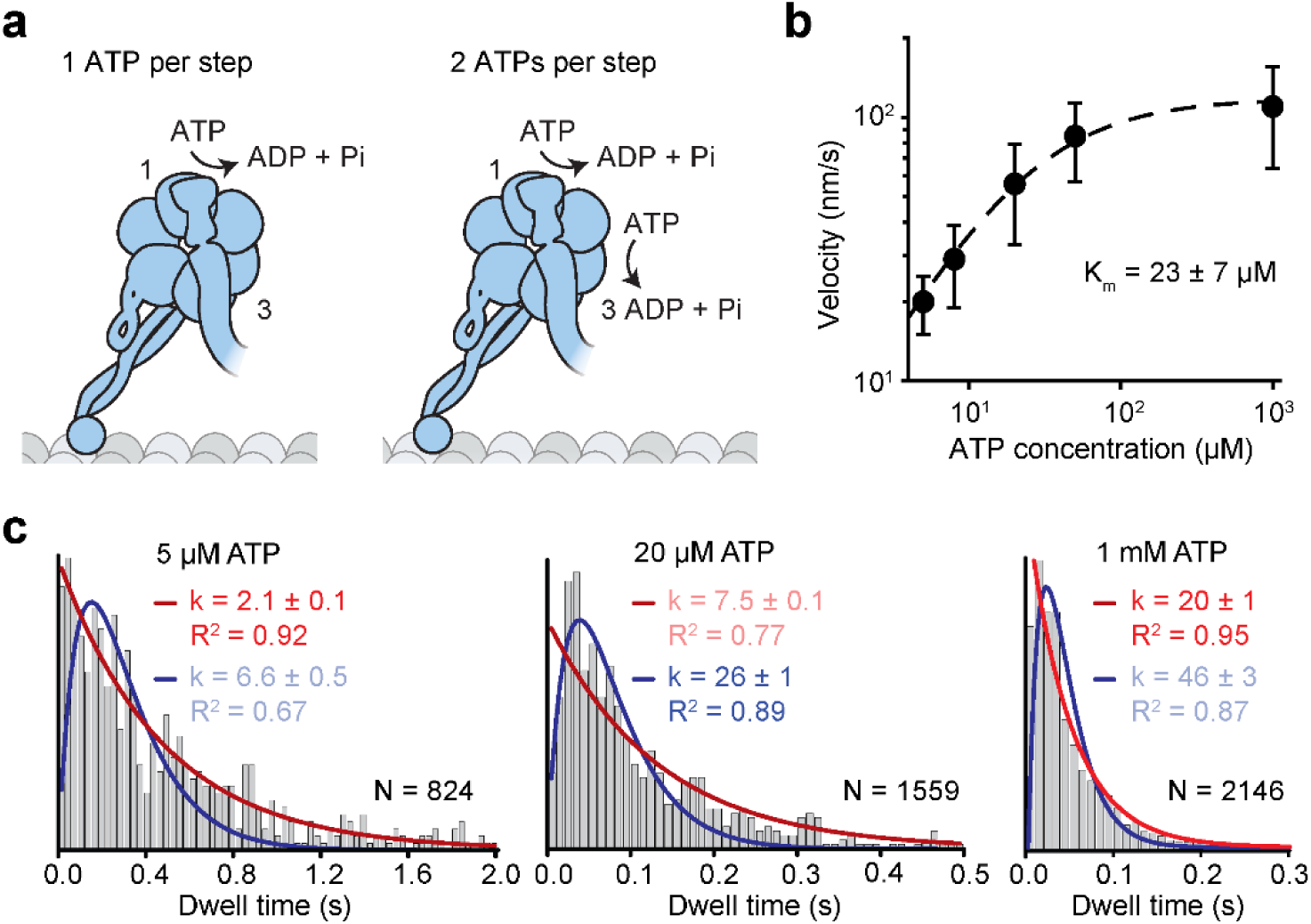
Dynein hydrolyzes one ATP per stepping cycle. **a,** Kinetic schemes of dynein hydrolyzing one or two ATPs per step. **b,** The mean velocity of analyzed trajectories under different ATP concentrations (mean ± s.d.; n = 14, 22, 22, 18, 20 motors from left to right). K_M_ is calculated from a fit to Michaelis-Menten kinetics (dashed curve). Error bars represent 95% c.i. **c,** The histogram of dwell times between successive steps of dynein along the MT long axis fits better to a single exponential decay (red curves) at limited (5 and 8 µM) and saturating (1 mM) ATP concentrations and to a Gamma distribution (blue curves) at K_M_ = 20 µM ATP. The parameters with a higher goodness of fit (R^2^) are opaque.

To determine how the two heads step relative to each other, we tracked the stepping of a dynein heterodimer labeled with different-colored fluorophores at each MTBD using two-color MINFLUX (Fig. 3a, Extended Data Fig. 6). As previously reported^4, 5^, dynein heads step independently of each other (Fig. 3a), but the stepping head tends to move towards the tethered head to minimize the intramolecular tension on the linkers that interconnect the two heads^4^ (Fig. 3b). As the inter-head distance increases along the MT long axis, the stepping rate of the motor increases and the trailing head becomes more likely to take a step (Fig. 3c-d, Extended Data Fig. 7), supporting the model that dynein takes tension-induced steps in addition to ATP-induced steps^36^. While the two heads frequently swap the lead position along the MT long axis, they were more likely to maintain left-right positions on the short axis (Fig. 3e, Video 1). These results indicate that dynein shuffles its two heads on separate protofilaments along the MT long axis^23^, while crossing of the heads in the MT short axis is more restricted by the tethered head.

**Figure 3.**
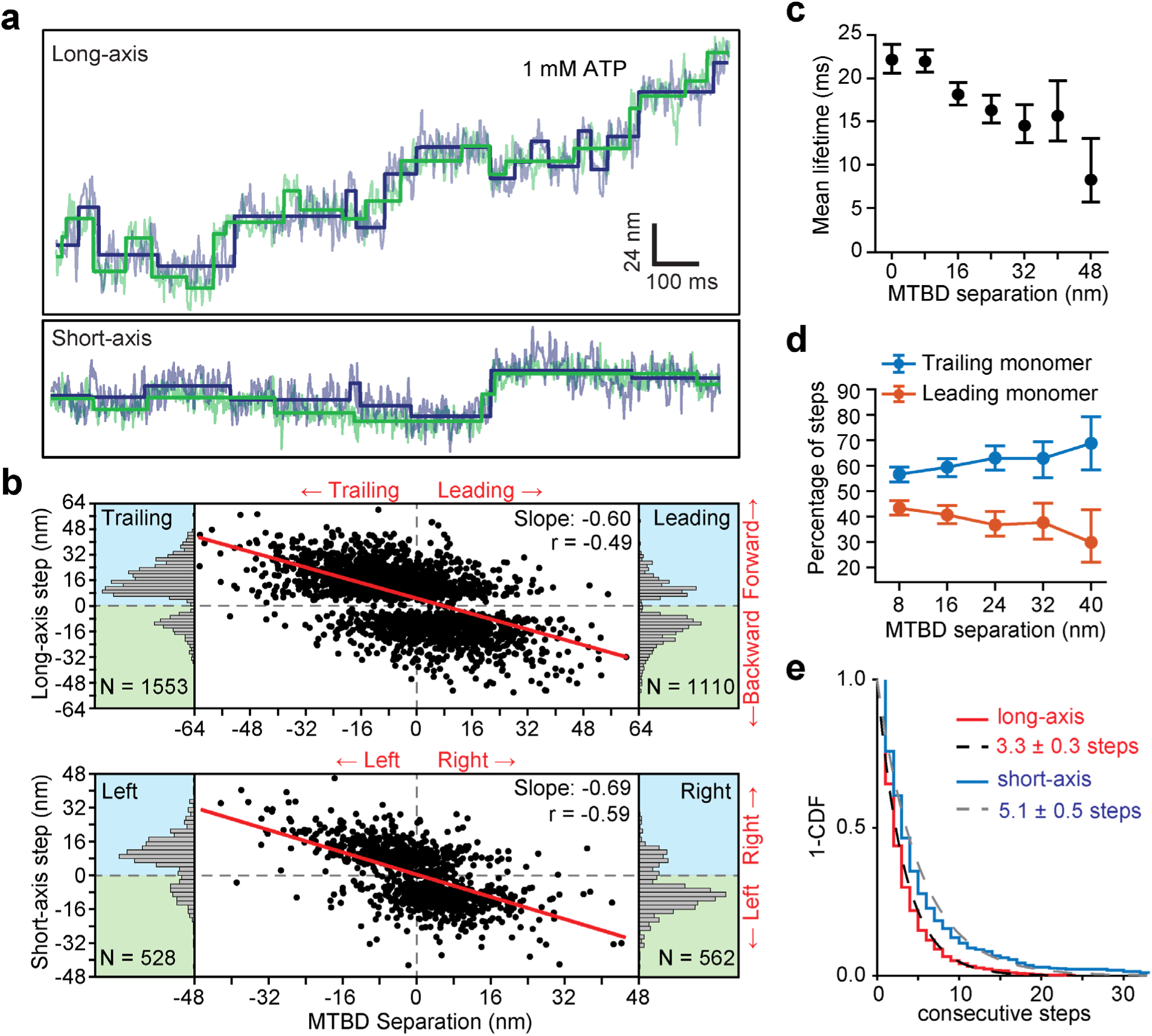
Two-color MINFLUX tracking of dynein at saturating ATP. **a,** Two-color MINFLUX trajectory of dynein dual labeled with LD555 (green) and LD655 (purple) tracked with a 3.3 nm and 640 µs resolution at 1 mM ATP and fit to a step-finding algorithm (solid lines). **b,** The size and direction of the steps taken along the MT long and short axes as a function of the relative position of the MTBDs. The red lines represent a linear fit (*r*: Pearson’s *r*). **c,** Mean lifetime of steps taken by dynein under different MTBD separations (n = 661, 1152, 730, 495, 170, 77, 23 steps from left to right). **d,** Percentage of steps taken by the leading or trailing head by the separation between their MTBDs (From left to right, n = 652, 433, 249, 106, 53 steps for the trailing and 500, 297, 146, 64, 24 steps for the leading head). **e,** The inverse cumulative distribution (1-CDF) of the stepping head was fit to a single exponential decay to calculate the average number of consecutive steps the head maintains its leading/trailing position on the long axis and right/left position on the short axis relative to the tethered head (n = 586 and 505 for long-and short-axes, respectively). In c and d, error bars represent 95% c.i.

In both one- and two-color MINFLUX trajectories, 14% of steps were preceded by a characteristic backward movement that typically lasts a single frame, referred to as dips (Fig. 1e, insert). Upon increasing the temporal resolution of MINFLUX to 0.28 ms, we detected a higher fraction (35%) of steps preceded by backward dips that varied from 8 to 24 nm in size (Fig. 4a-c). The lifetimes of the dips were independent of ATP concentration and much shorter (5-7 ms) than those of the steps (∼40 ms in saturating ATP; Fig. 4d, Extended Data Fig. 8), indicating that dips are not backward steps. Instead, they represent a transient intermediate that occurs after ATP binding and before a head takes a step. Two-color trajectories revealed that both the leading and trailing heads exhibit dips with nearly equal probability before taking a step (Extended Data Fig. 8f), ruling out the possibility that backward dipping occurs only when the leading head releases from the MT and is pulled back toward the trailing head. The transient nature and variable size of the dips provide an explanation for why we observed only a third of the steps preceded by a clearly distinguishable dip.

**Figure 4.**
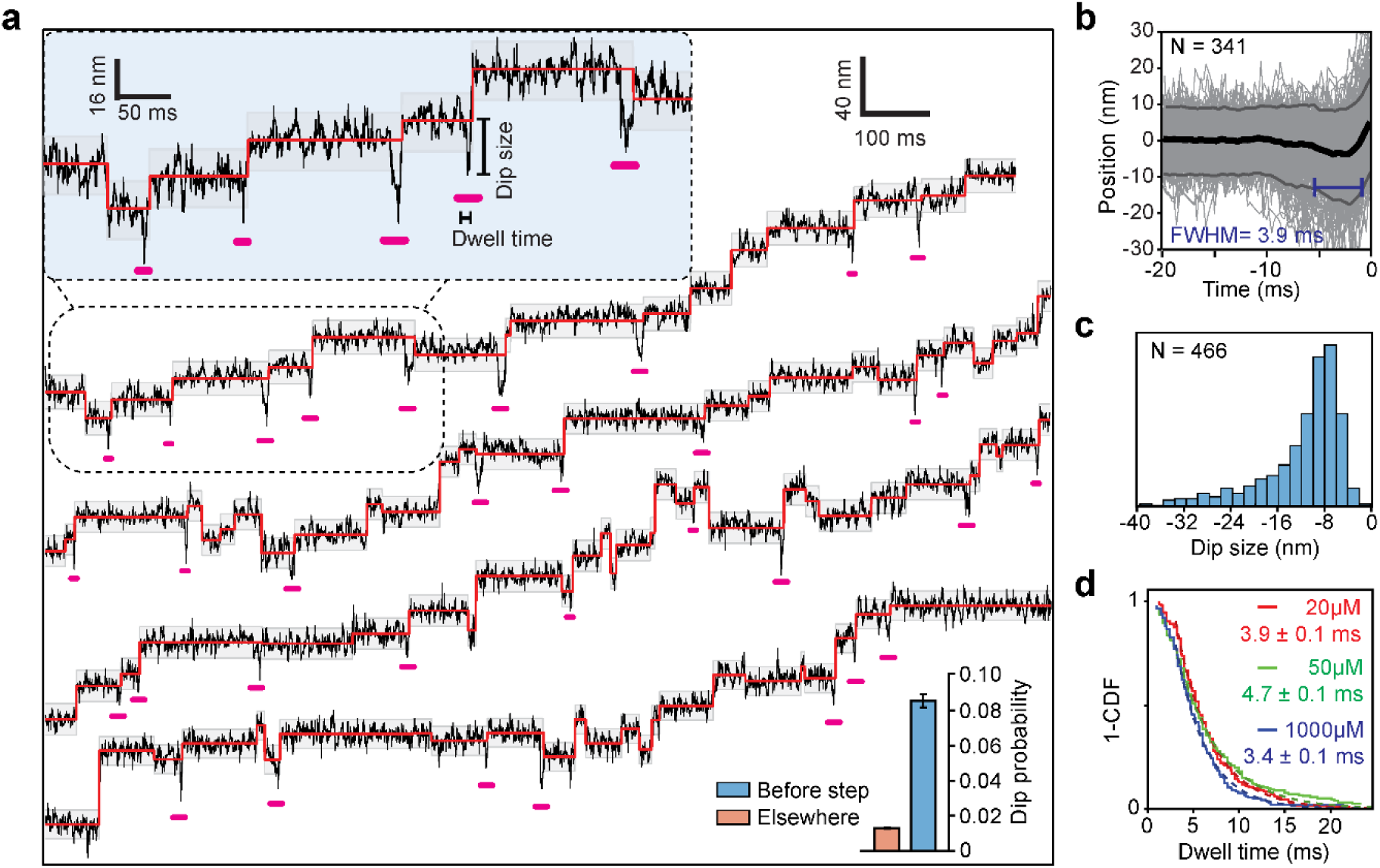
Dynein steps are preceded by backward dips. **a,** Example trajectories of dynein in 1 mM ATP at 280 µs temporal resolution show brief dips (underlined in pink) before the steps. Shaded regions represent ±2σ of the dwelling position between successive steps. (Bottom insert) The probability of extended -2σ deviation from the dwelling position within 7 ms before a step or elsewhere in the trajectories (±95% c.i.). **b,** The cumulative overlay of the dwell positions before a step (t = 0 s) demonstrates the presence of a backward dip. Black and grey curves represent mean ± 2σ, respectively. The average duration of the dips was measured from the full-width half minimum (FWHM) of the -2σ deviation before steps with dwell times larger than 50 ms. **c,** The histogram of the dip size. **d,** 1-CDF of dip lifetimes recorded in 2.8 ms temporal resolution at different ATP concentrations. The lifetime of dips (±s.e.) was calculated by a fit to a single exponential decay (solid curves; n = 280, 193, 322 from top to bottom).

We reasoned that dips occur upon the release of the stepping head from the MT after it binds ATP (Extended Data Fig. 9). Because the dynein motor domain is tilted relative to the MTBD^20^, the link between the two heads sits toward the plus end of both MTBDs. Therefore, upon MT dissociation, the MTBD of the stepping head moves backward and diffuses around the tethered head until dynein undergoes a conformational change that moves it forward. According to this scheme, the end of a dip marks when dynein generates forward bias in its stepping cycle. To determine which chemical process in the ATPase cycle terminates the dip, we slowed ATP hydrolysis by mutating the AAA1 site (Fig. 5a). A well-conserved aspartate (D) residue in the Walker B motif coordinates the Mg^2+^ ion for proper ATP hydrolysis^19, 37^, and replacing it with a bulkier glutamate (E) slows ATP hydrolysis in other AAA enzymes^38^. Consistently, molecular dynamics (MD) simulations showed that the D1848E mutation at AAA1 of dynein shifts the Mg^2+^ ion away from the β- and γ-phosphate oxygens, suggesting slower ATP hydrolysis (Fig. 5a, Extended Data Fig. 10, Video 2). The D1848E mutant had similar velocity, run length, and stepping properties to WT dynein (Fig. 5b, Extended Data Fig. 11). Remarkably, D1848E has a two-fold increase in dip probability compared to WT dynein (Fig. 5c). In comparison, the addition of excess (30 mM) inorganic phosphate to stabilize the ADP.Pi state^39, 40^ of dynein does not affect the motility and the dipping behavior (Fig. 5b-c, Extended Data Fig. 11). These results are consistent with a scheme where dips terminate after ATP hydrolysis and before phosphate release (Fig. 5d), indicating that dynein generates forward bias upon ATP hydrolysis.

**Figure 5.**
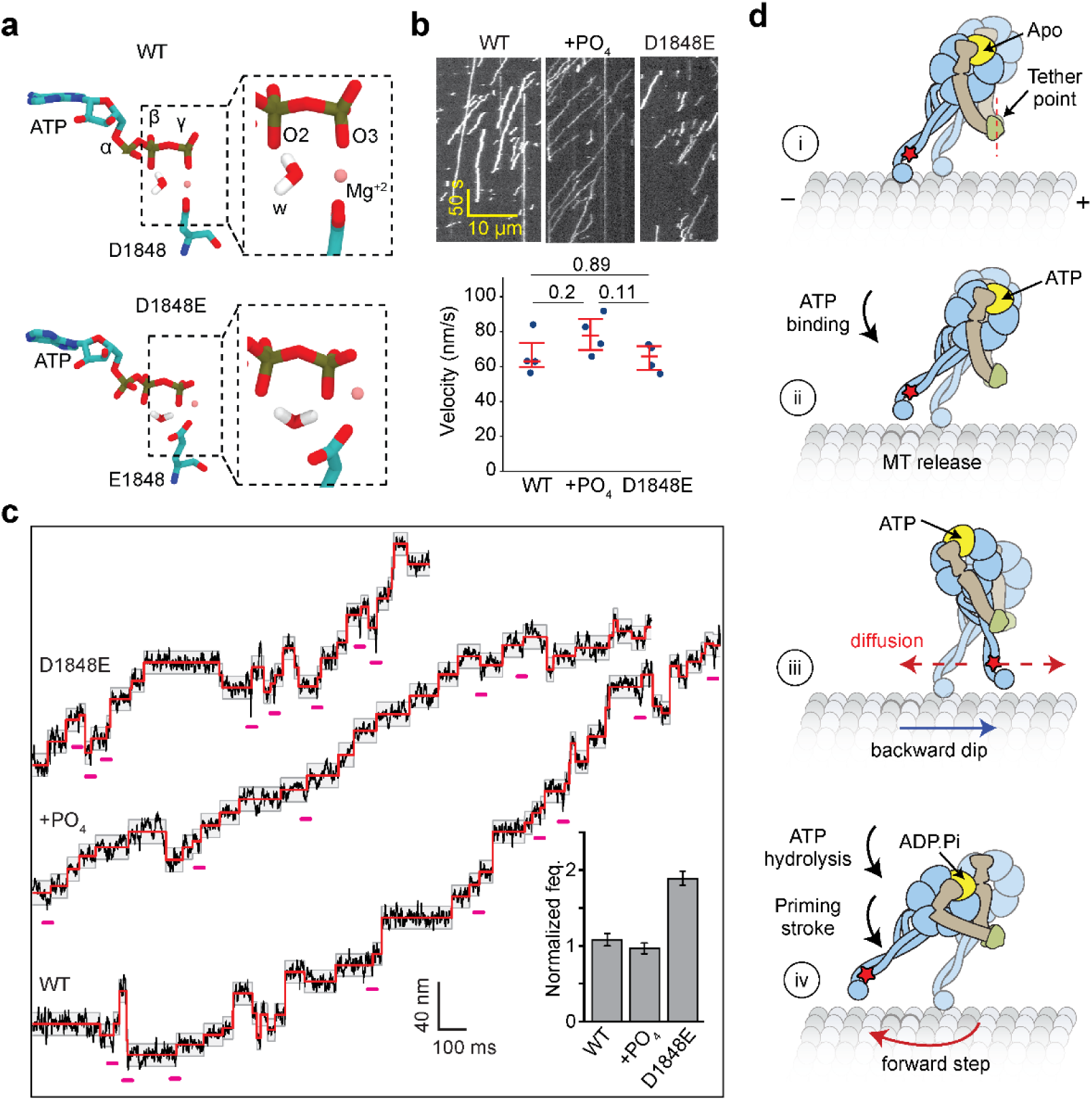
Dynein generates a net forward bias upon ATP hydrolysis. **a,** MD simulations show that Mg^2+^ and water (w) coordinate the β- and γ-phosphate oxygens (O2 and O3) of ATP in WT dynein. In D1848E, Mg^2+^ is shifted away from its canonical site, with O3 positioned between Mg²⁺ and O2, effectively blocking their coordination. **b,** Example kymographs (top) and average velocities (bottom) of WT dynein, WT with 30 mM PO_4_ (+PO_4_), and D1848E at 1 mM ATP. The centerline and whiskers represent mean and 95% c.i., respectively (n = 4 biological replicates). *P* values were calculated by the Mann-Whitney U-test. **c,** Example trajectories of WT and D1848E trajectories tracked with MINFLUX at 3.6 nm and 0.28 ms resolution. Detected dips are underscored in pink. (Insert) The normalized frequency of detecting a dip before a step (mean ± 95% c.i.). **d,** Interpretation of the dips observed in MINFLUX trajectories. (i) Dynein is strongly bound to the MT in the nucleotide-free state (Apo). (ii) Upon ATP binding, a dynein head releases from the MT and (iii) moves backward toward the tethering point connecting the heads before ATP is hydrolyzed. (iv) After ATP hydrolysis (ADP.Pi), the linker undergoes the priming stroke and moves the MTBD towards the minus end before dynein rebinds the MT.

## Discussion

Our MTBD labeling combined with MINFLUX resolves the sequence of dynein’s mechanochemical events at millisecond and nanometer scales, which cannot be captured by cryo-EM. Directly labeling the MT binding interface enabled a clear assignment of MT release, diffusional search, and productive rebinding of dynein. The stepping statistics across ATP concentrations, along with the analysis of stepping kinetics, support a one-ATP-per-step mechanism. The observed 8-nm periodicity with large variance, frequent access to neighboring protofilaments, and a measurable fraction of apparent 0-nm steps characterize a search-and-capture process that is naturally biased toward the minus end but remains highly variable at the single-step level.

Two-color MTBD tracking further refines the inter-head coordination of dynein. The heads step mostly independently, yet intramolecular tension increases the probability that the trailing head steps as the inter-head distance grows in the MT long axis^36^. These results are consistent with faster MT detachment of the trailing head under forward tension, whereas the leading head resists detachment under backward tension^41^. The stepping rate of dynein also increases with the interhead separation along the short axis (Extended Data Fig. 7d-e), but we did not observe differences in the stepping probabilities of the right and left heads, indicating that dynein does not exhibit asymmetric detachment kinetics across the perpendicular axis of MTs. Long-axis positions swap often, but short-axis positions stay constant for nearly two times long, implying that the MT tethered head discourages head crossover across protofilaments. These rules explain how dynein can combine a powerstroke mechanism and stochastic head diffusion to sustain unidirectional motility with highly variable step size and occasional backsteps. Diffusion and linker geometry sometimes return the head to the original binding site before the cycle commits, resulting in foot stomping of dynein.

The short, backward “dip” that occurs immediately before steps indicates a transient, ATP-binding–induced release state followed by relaxation of the detached head toward the dimer’s tether point. The analysis of the mutant dynein trajectories indicates that the detached head diffuses around the tether point until it hydrolyzes the ATP. Upon ATP hydrolysis, the linker of the detached head switches from a straight conformation to semi-bent and bent conformations across the ring, or partially detaches from the ring^19, 21, 42, 43^. This priming stroke pushes forward the MTBD of the detached head, which rebinds the MT after phosphate release (Extended Data Fig. 9b). Diffusional search of the detached head and different conformations of the linker may contribute to the high variability in the dynein step size (Fig. 5d, Extended Data Fig. 9c). Alternatively, the priming stroke only increases the distance between the tether point and the MTBD of the detached head without pushing it forward. This conformational change effectively increases the radius of diffusion for a new tubulin binding site, and the detached head becomes more likely to bind forward because of the requirement of the angle of the stalk pointing forward^20^. However, this latter model would predict that the detached head continues to diffuse around the tether point in the ADP.Pi state, which is inconsistent with the lack of increase in dipping in the presence of excess phosphate.

The ability to site-specifically label Dyn_CLM_ also opens new ways of visualizing the coordinated action of the AAA ring, stalk, and linker during dynein stepping in real time. MINFLUX provides the time resolution required to observe conformational dynamics of the linker in a single ATPase cycle. The combination of MINFLUX with single-molecule FRET^44^ can directly observe how conformational changes of the linker couple to the MT attachment/detachment and forward motion to provide a more complete picture of the mechanochemical cycle.

## Acknowledgements

We thank M. DeWitt and A. Yonar for preliminary work on FIONA, J. Matthias, J.O. Wirth, K. Boateng, and G. Fried for technical assistance with the MINFLUX microscope in IGB at the University of Illinois. This work was funded by grants from the NSF Science and Technology Center for Quantitative Cell Biology (2243257 to P.R.S.), NSF (MCB-1055017 and MCB-1617028 to A.Y., and DGE 2146752 to J.S.), NIH (GM136414 to A.Y.; GM132392 to P.R.S.), and the Medical Research Council (MC_UP_A025_1011 to A.P.C.).

## Author contributions

J.S., E.G-H., A.P.C., P.R.S., and A.Y. conceived the study and designed the experiments. E.G-H., A.P.C., and J.S. created the constructs. J.S. and E.G. prepared, isolated, and labeled the proteins. J.T.C. performed FIONA experiments. J.S. and D.P.W. performed MINFLUX experiments. M.Gur and M. Golcuk performed MD simulations. J.S., P.R.S., and A.Y. wrote the manuscript, and all authors read and commented on the manuscript.

## Competing Interests

The authors declare no competing interests.

## Materials and Methods

## Cloning and molecular biology

*S. cerevisiae* cytoplasmic dynein heavy chain gene (*DYN1*) was truncated at the N terminus (encoding amino acids 1219–4093, referred to as Dyn_314kDa_) as a template for mutagenesis^23^. A tandem protein-A (ZZ) tag, a dual Tev protease site, and a glutathione-S-transferase (GST) dimerization tag were added to the N-terminus of the Dyn_314kDa_ gene, and the entire construct was cloned into the ampicillin and URA3-selectable pYES2 plasmid downstream of the inducible galactose promoter. The cysteine-light dynein construct was generated by stitching PCR and Gibson assembly. For heterodimerization, the GST tag was replaced by SpyTag and SpyCatcher. Plasmids were transformed into the CY1 yeast acceptor strain (W303 MATa; his3-11,15; ura3-1; leu2-3,112; ade2-1; trp1-1) using lithium acetate (LiAc) transformation to enable exogenous expression in synthetic defined media plates lacking uracil (SD Ura). The list of plasmids and yeast strains used in this study is given in Extended Data Table 3.

The crystal structure of the *S. cerevisiae* dynein motor domain (PDB ID 4AKG)^27^ was used to analyze the accessibility of cysteine residues by PyMOL. Cysteine residues with more than 5 Å, between 2.5 - 5 Å, and less than 2.5 Å surface accessibility were classified as surface exposed, partially exposed, and buried, respectively (Extended Data Table 1). The five surface-exposed residues on the dynein motor domain were mutated to serine (Dyn_CLM_). Partially exposed cysteines of dynein could not be removed as their mutagenesis substantially lowered protein expression (not shown). Surface-exposed cysteines of the GST-tag were also mutated to serine.

### Protein expression, purification, and labeling

Dynein proteins were expressed in yeast. We found that expression could be performed in YP media without selection for the pYES plasmid. A single fresh yeast colony was used to inoculate 10 mL of YP media containing 2% glucose. Cultures were grown overnight at 30°C with shaking at 200 rpm. The 100 ml YP media containing 1% raffinose was inoculated with an overnight culture for ∼8 h at 30°C at 200 rpm agitation until OD_600_ reached 0.2. The 100 ml culture was used to inoculate 2 L of YP media containing 2% galactose and 100 mg/ml adenine (Sigma) for 24-48 h at 30°C with 200 rpm agitation. Yeast was harvested by centrifugation at 5000 *g* for 7 min. Pellets were resuspended in phosphate-buffered saline to form a thick paste and frozen by dropwise addition in liquid nitrogen. Frozen yeast was stored at -80°C.

Frozen yeast pellets were ground and thawed in a lysis buffer (30 mM HEPES, pH 7.4, 50 mM K-acetate, 2 mM Mg-acetate, 1 mM EGTA, 0.1 mM ATP-Mg^2+^, 1 mM DTT, 2 mM PMSF). The lysate was mixed with IgG Sepharose affinity beads and then cleaved by TEV protease, as described previously^23^. The resulting protein was concentrated and stored in storage buffer (30 mM HEPES, pH 7.4, 175 mM KAc, 2 mM MgAc_2_, 1 mM EGTA, 10% glycerol, 0.2 mM TCEP, 0.1 mM ATP). The proteins were run in a denaturing gel, and their concentrations were determined from OD_280_.

20 pmol SpyTag dynein was then incubated with 30-fold excess maleimide reactive dyes in TEV storage buffer at room temperature for 1 h. The reaction was quenched with 1 mM DTT. After removing excess dye with a Zeba desalting column, SpyCatcher-dynein was incubated with SpyTag-dynein at a 1:5 ratio at room temperature for 20 min to form heterodimers. Dimerization was confirmed using native gel and size exclusion chromatography. Fluorescence labeling was detected using a Typhoon gel image scanner. The labeling efficiency (∼80%) was determined from 280 nm and 561 nm absorbance in a Nanodrop spectrophotometer. Purified protein was aliquoted, flash-frozen in liquid nitrogen, and stored at -80°C.

### MT polymerization

Tubulin was purified from pig brains in a 1 M PIPES buffer. 120 ng unlabeled, 25 ng biotinylated, and 15 ng fluorescently-labeled tubulin were mixed in BRB80 buffer (80 mM PIPES, pH 6.8, 2 mM MgCl_2_, 1 mM EGTA) supplemented with 1 mM GTP and DTT at 4°C. The 20 µL mixture was cleared by incubating on ice for 10 min and then centrifugation at 290,000 g for 10 min at 4°C. The supernatant was transferred to a 37 °C bath. Taxol was serially added to the final concentration of 0.1 µM, 1 µM, and 10 µM following 10 min incubation periods. After the final polymerization step of 15 min, the taxol-stabilized MTs were spun at 21,000 g for 15 min at 37°C to pellet polymerized MTs. The pellet was then resuspended in 20 µL pre-warmed BRB80 buffer supplemented with 1 mM DTT and 10 µM taxol. MTs were stored in the dark at room temperature and used within two weeks.

### Single-molecule motility assays

Motility assays were performed in custom-made flow chambers comprised of polyethylene glycol (PEG)/PEG-biotin-coated coverslips adhered to glass slides by double-sided tape^45^. The chamber was incubated with 10 µL 0.5 mg/mL streptavidin and washed with 40 µL DLB-CPT buffer (30 mM HEPES, pH 7.2, 20mM KAc, 2 mM MgCl_2_, 1 mM EGTA, 10% glycerol with added 1 mg/mL casein, 0.5% pluronic acid, and 1 µM Taxol). The chamber was then incubated with Cy3- and biotin-labeled MTs for 3 min before washing with 60 µL DLB-CPT. Dynein was then diluted in DLB-CPT and flown into the chamber. Motor concentration was adjusted to ensure fewer than one fluorescently labeled motor run per µm length of the MT during imaging. After 3 min of incubation, the unbound motor was washed with 10 µL stepping buffer (SB) containing DLB-CPT, 0.8% dextrose, 0.1 mg/mL glucose oxidase, 0.2 mg/mL catalase, and the desired ATP concentration. The sample was sealed and imaged for 1 h. Two-color MINFLUX and one-color MINFLUX GST-dynein experiments were performed using Atto488- and biotin-labeled MTs, and 0.1% methylcellulose was added to the final SB solution.

### Microscope and imaging

Single-molecule motility assays were performed on a custom-built objective-type total internal reflection fluorescence (TIRF) microscope, equipped with an inverted Nikon Ti-E microscope body, a perfect focusing system, and a 1.49 NA 100× oil immersion objective. The sample was illuminated with 488, 561, and 633 nm laser beams (Coherent) to excite GFP/Atto488, Cy3/Alexa555/LD555, and Cy5/Alexa647/LD655 fluorophores, respectively. Movies of LD655-labeled dynein were recorded with a 0.1 – 0.3 s exposure time under 2 mW 633 nm excitation. The fluorescence signal was detected with an electron-multiplying CCD camera (Andor, iXon).

Single-molecule tracking measurements were also made on a commercial MINFLUX instrument (Abberior). The microscope is equipped with a 100x 1.4 NA oil immersion objective lens (Olympus), 561 nm and 642 nm excitation lasers, avalanche photodiodes (Excelitas) with a detection range to capture LD55 and LD655 fluorescence, and a pinhole size corresponding to 0.78 airy units. The microscope was controlled by Abberior Imspector. An iterative localization approach with a hexagonal array of points was used to localize the fluorophores, as reported previously^30, 46^. Once the fluorophore was detected, the radius of the donut-shaped focused excitation beam (L) was decreased (284 nm, 302 nm, 151 nm, and 75 nm) and the laser power was increased (60, 60, 120, and 150 µW) in three increments. At this final step, ∼100 photons were collected before localization. In two-color MINFLUX assays, the excitation lasers were interleaved between valid localizations. This final iteration was repeated continuously until the detection signal was lost due to permanent photobleaching of the dyes. This procedure achieves ∼2.5 ms for tracking of the LD655 dye. The parameters of the tracking algorithm were adjusted to increase the temporal resolution (Extended Data Table 4).

### Alignment of Two-Color Trajectories

Both 561 and 642 nm toroidal excitation patterns were properly shaped and aligned at diffraction-limited resolution with minimal differences in the angular intensity. Because the z-position of the microscope objective significantly changes the registered position of the two channels, we performed local aberration correction by collecting two-color trajectories in the same 6 µm x 2 µm area. The mean offset in the x-y plane for each temporally correlated pair of points of both channels was calculated for all trajectories that contain at least 300 data points. Outlier trajectories more than 30 nm away from the mean were excluded. The mean offset in *x* and *y* directions was adjusted as a linear correction to align the two channels (Extended Data Fig. 6).

### Data processing and analysis

Recorded TIRF movies were analyzed by a two-dimensional Gaussian fitting algorithm on a custom MATLAB software, YFIESTA (https://github.com/Yildiz-Lab/YFIESTA), to localize the fluorescent spots in the *xy* plane. All MINFLUX data were also processed, analyzed, and rendered on YFIESTA. Trajectories longer than 100 nm were detected and sorted according to a custom algorithm that relies on time autocorrelation along a linear path to determine fitness. The *x* and *y* axes of the trajectories were rotated accordingly to match the long and short axes of the MT tracks. Trajectories that had an acquisition frequency greater than 100 kHz (suggesting multiple photon emissions), high noise in the short axis due to poor MT immobilization, and diffusive or poor motility were excluded from data analysis. Trajectories that passed initial screening were then fit to steps by a Schwartz Information Criterion-based step-fitting algorithm in both the long- and short-axis of MTs, as described previously^41^. Steps that were less than 3 frames were removed. All steps were subject to a windowed mean filter. If the difference of the mean of 8 data points before and after a step was less than 7.5 nm, these steps were removed to eliminate the positional drift being detected as a step. Step fitting was then visually inspected and manually corrected to prevent overfitting of the trajectories. In a two-dimensional step analysis, long-axis and short-axis steps that occur within 5 frames (∼12 ms) were combined into a “diagonal” step.

Analysis of dips was performed on the long axis of the trajectories. A custom algorithm detects the deviation of a data point more than two standard deviations (2σ) of the mean dwelling position before a detectable step. Dips were defined as 2σ deviations within 3 (∼7.2 ms) or 20 data points (∼7 ms) before a detectable step for trajectories recorded at ∼2.5 or 0.28 ms temporal resolution, respectively. The dwell time of a dip was defined as the time between the beginning of a dip and the movement of the stepping head to the next position. The statistical significance of dips was calculated by comparing the probability of observing a -2σ deviation within ∼7.2 ms (3 and 20 data points at ∼2.5 and ∼0.28 ms temporal resolution, respectively) around a step versus anywhere in the trajectory. The Beta distribution confirmed with >95% confidence that -2σ deviations are not randomly distributed, and instead frequently occur before steps.

The two channels in 2-color MINFLUX data were analyzed separately for step fitting and then registered to determine the interhead separation vector before and after either head takes a step. Concurrent stepping positions were calculated by determining the number of steps a head maintains its relative position to its partner in the long- or short-axis of MTs. The heads were assumed to maintain their position if they were within 3 nm of each other on the analyzed axis after a step. The leading/trailing and left/right positions of the heads were recorded in a sequence after each step, and the 1-CDF of this relative position in sequence was fit to determine the average number of steps that a head maintains its relative position.

### MD Simulations

The MT-bound structure of dynein was constructed using the cryo-EM structure of the motor domain of human cytoplasmic dynein-1 (PDB ID 7Z8F; residues Q1327-E4646), which provides backbone coordinates but lacks side-chain information. The conformations for the AAA+/catalytic ring (residues Q1327*–D3221 and A3470-E4646*) were adopted from PDB ID 7Z8G, and MTBD side-chain conformations (residues A3272*C–E3420*) were modeled using PBD ID 6RZB. The remaining stalk (L3222-I3271 and D3421-E3469) side-chain conformations were modeled in VMD using the CHARMM36 force field. To obtain the ATP-bound state of the AAA1 domain, ADP at AAA1 was replaced with ATP, and the ATP/Mg²⁺ coordinates were taken from the AAA2 site. The D1958E mutation (corresponding to D1848E in yeast dynein) was introduced using the VMD Mutator plugin. Both WT and mutant systems were solvated with at least 15 Å of water padding in each direction. Ions were added to neutralize the system and to set the ionic concentrations to 1 mM MgCl_2_ and 150 mM KCl. The solvated and ionized system is composed of approximately one million atoms.

All MD simulations were performed using NAMD 3 with the CHARMM36m all-atom additive protein force field and a time step of 2 fs. A 12 Å cutoff distance was applied for van der Waals interactions, and long-range electrostatics were calculated using the particle–mesh Ewald method. The temperature was maintained at 310 K using Langevin dynamics thermostat with a damping coefficient of 1 ps⁻¹, while the pressure was controlled at 1 atm using the Langevin piston Nosé–Hoover barostat with an oscillation period of 100 fs and a damping constant of 50 fs. Each system was first energy-minimized for 10,000 steps, followed by 2 ns of minimization with proteins, nucleotides, and bound ions fixed. Subsequently, all restraints were released, and the systems underwent an additional 10,000-step minimization.

Equilibration was then performed in stages: first, with C_α_ atoms constrained using a 1 kcal/mol·Å^2^ force constant for 5 ns, followed by 5 ns of equilibration, then production runs with tubulin atoms constrained with a spring constant of 1 kcal/mol·Å^2^ to mimic protofilament movements. For production runs, WT systems were simulated in two independent sets of 1 μs each, yielding a total of 2 μs. Mutant systems were simulated in four independent sets of 300 ns each, yielding a total of 1.2 μs.

## EXTENDED DATA FILES

**Extended Data Figure 1.**
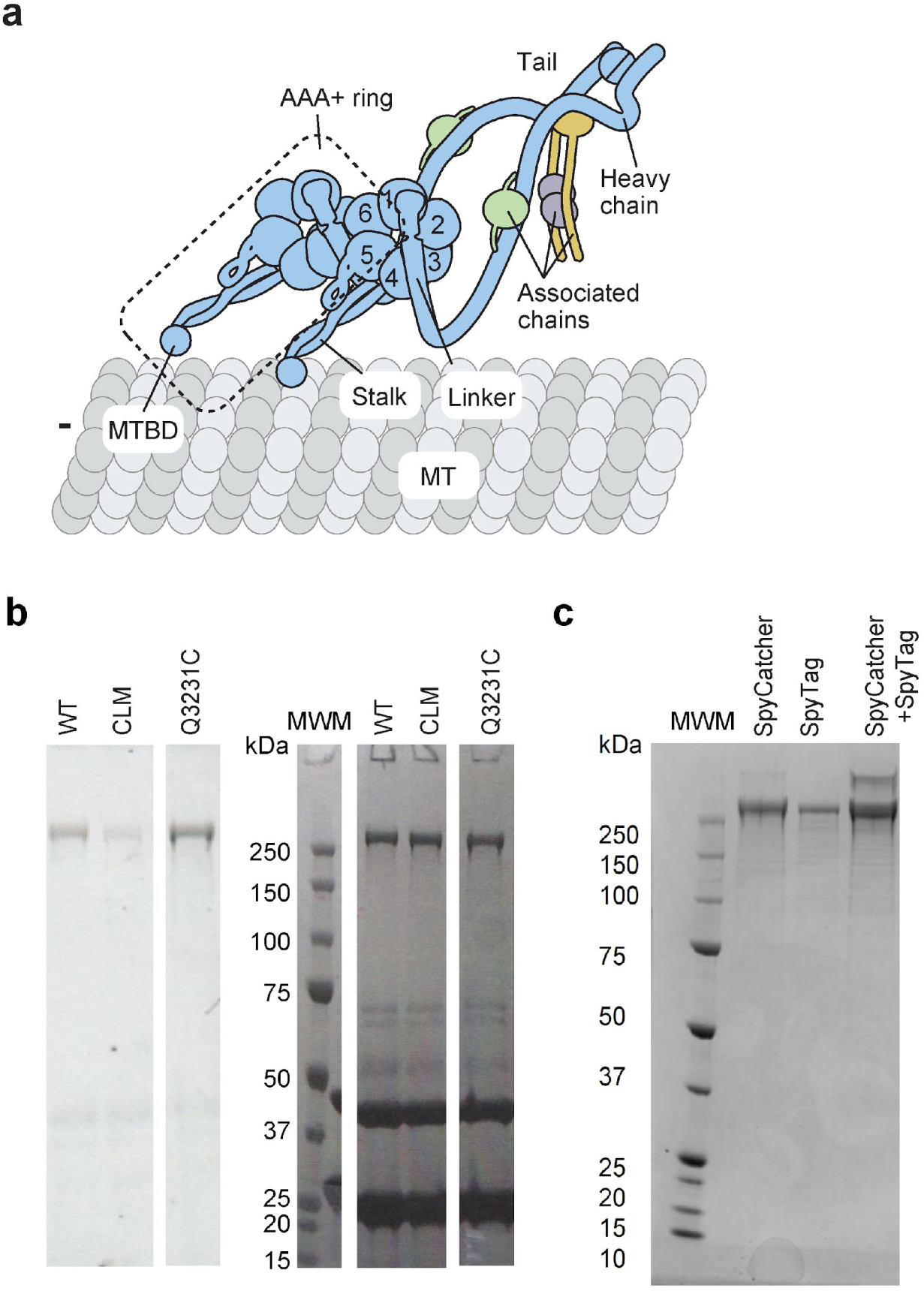
Labeling and heterodimerization of the Dyn_CLM_ mutant. **a,** The schematic of yeast dynein dimer bound to the MT. The dashed rectangle shows the motor domain of the dynein heavy chain. **b,** (Left) Fluorescence imaging and (Right) denaturing gel picture of GST-dynein constructs labeled with Alexa647-maleimide under the same experimental conditions (MWM: molecular weight marker; kDa: kilodalton). The bands in 27 and 43 kDa correspond to Tev protease and maltose binding protein (MBP) from the Tev-MBP construct used to elute the protein from affinity beads during preparation. **c,** A native gel picture shows the mixing of SpyCatcher-Dyn_CLM_ and SpyTag-Dyn_CLM_ at a 5:1 ratio to result in the heterodimerization of these constructs.

**Extended Data Figure 2.**
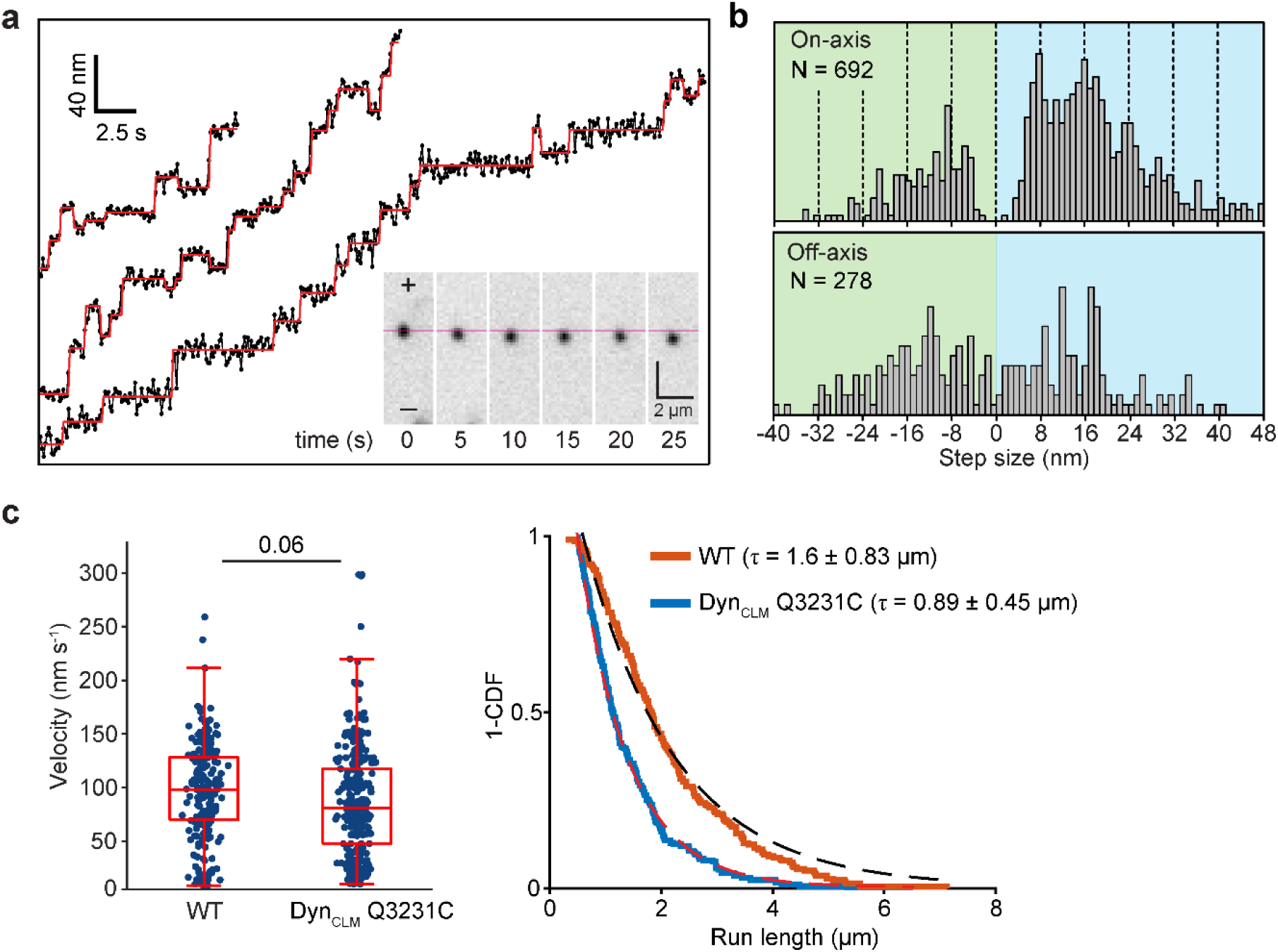
FIONA tracking of Dyn_CLM_ Q3231C. **a,** (Inset) Diffraction-limited images of a dynein heterodimer labeled with a single LD655 dye at its MTBD. Trajectories show FIONA tracking of dynein heterodimers with 100 ms temporal resolution at 2 µM ATP. The red lines show a fit for the step-finding algorithm. **b,** The histogram of steps taken along the MT long- and short-axis. Dashed lines show increments of 8 nm steps. **c,** Velocity and run length characteristics at saturating ATP of WT dynein and Dyn_CLM_ Q3231C. The centerline and whiskers represent the mean and 95% c.i., respectively (n = 218 for WT and 281 for Dyn_CLM_ Q3231C). P-values are calculated by a two-tailed t-test.

**Extended Data Figure 3.**
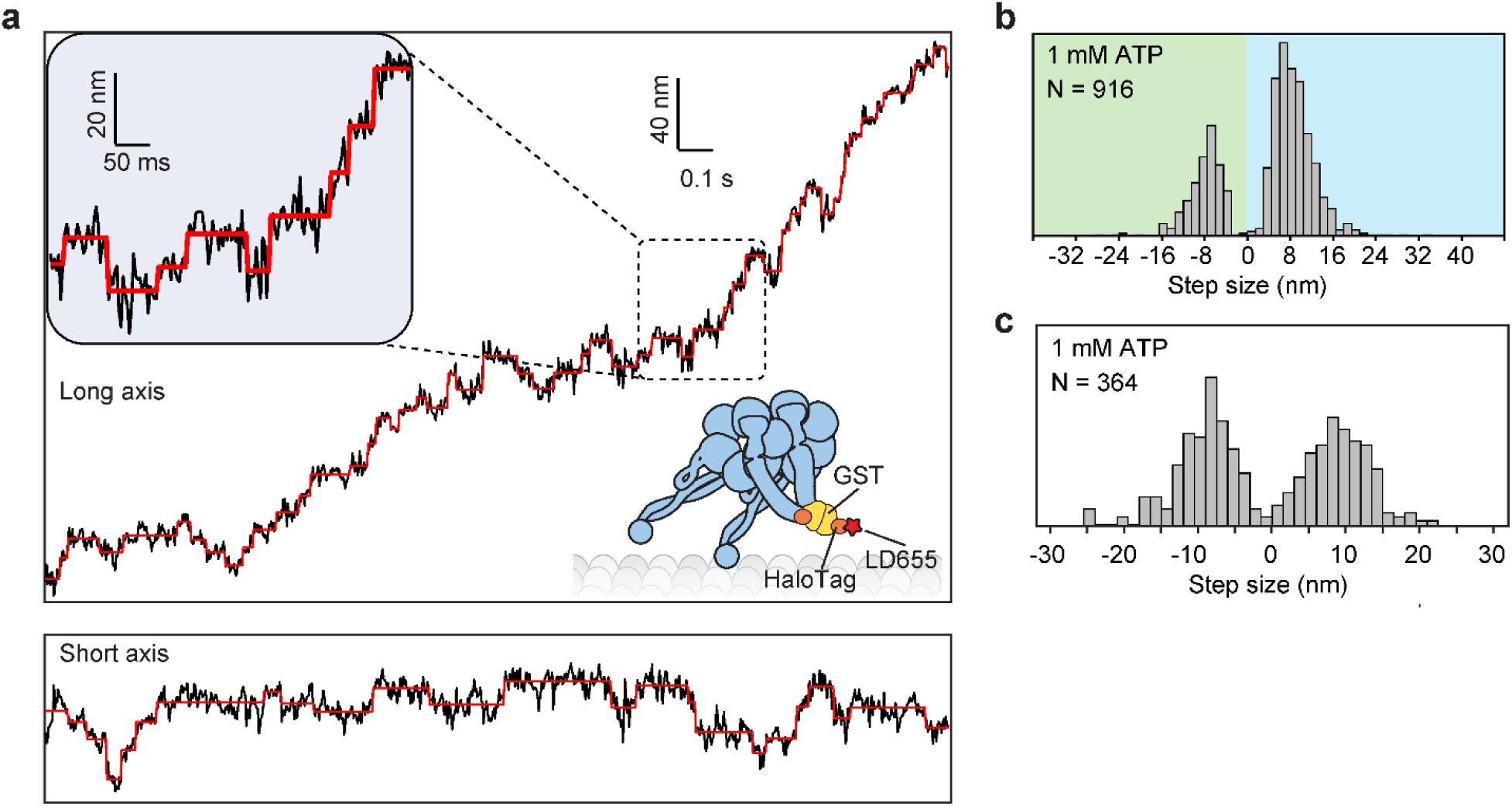
MINFLUX analysis of tail-labeled dynein. **a,** An example trajectory of GST-Dyn_314kDa_ labeled at its tail domain. The assay was performed at 1 mM ATP. Long- and short-axis displacement of the same motor are shown separately. Red horizontal lines represent a fit to the step finder algorithm. Despite high precision in localization (σ = 3.2 nm), the insert shows large fluctuations in the dye position while the motor dwells on the MT between steps. **b,** The histogram of the size steps taken by dynein in the long axis of MTs. **c,** The histogram of the size steps taken by dynein in the short axis of MTs.

**Extended Data Figure 4.**
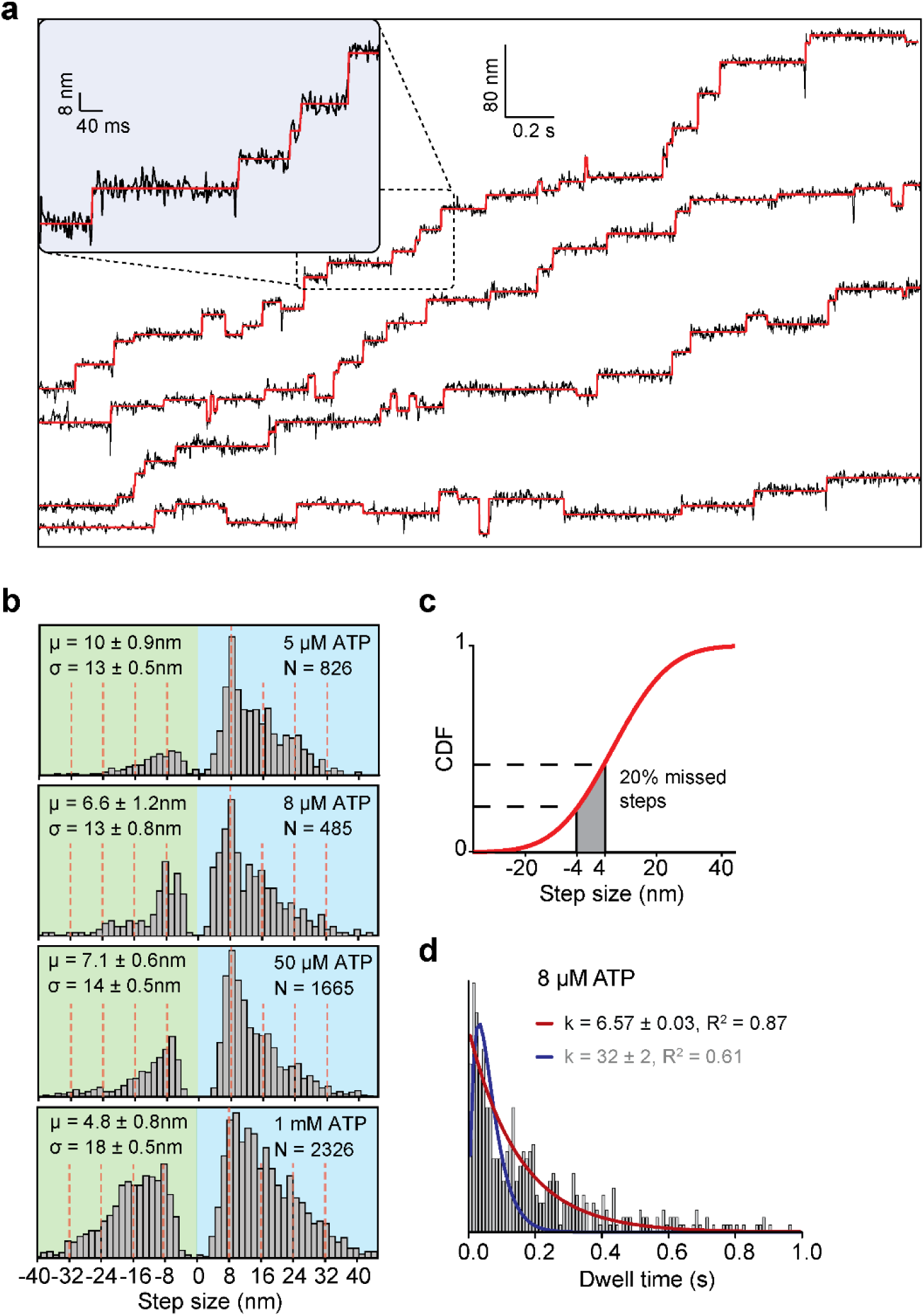
The step size distribution suggests “0 nm” steps. **a,** Additional example trajectories of dynein stepping along the MT long-axis at 20 µM ATP. **b,** Long-axis step size histograms under different ATP concentrations. Red dashed lines show increments of 8 nm steps ranging from -24 nm to 40 nm. Dynein has a similar step size distribution across 8 µM to 1 mM ATP (p >0.2, the Mann–Whitney U-test). **c,** The cumulative distribution function (CDF) of the Gaussian fit to Fig. 1g. CDF values from -4 nm to 4 nm suggest that 20% of the steps are 0 nm in size due to the rebinding of dynein to the same tubulin site (n = 5972 steps). **d,** The dwell time distribution of dynein stepping fits better to a single exponential decay (red curve) than a Gamma distribution (blue curve) at limited (8 µM) ATP. The fit parameters with a higher R^2^ are opaque. (n = 471 steps).

**Extended Data Figure 5.**
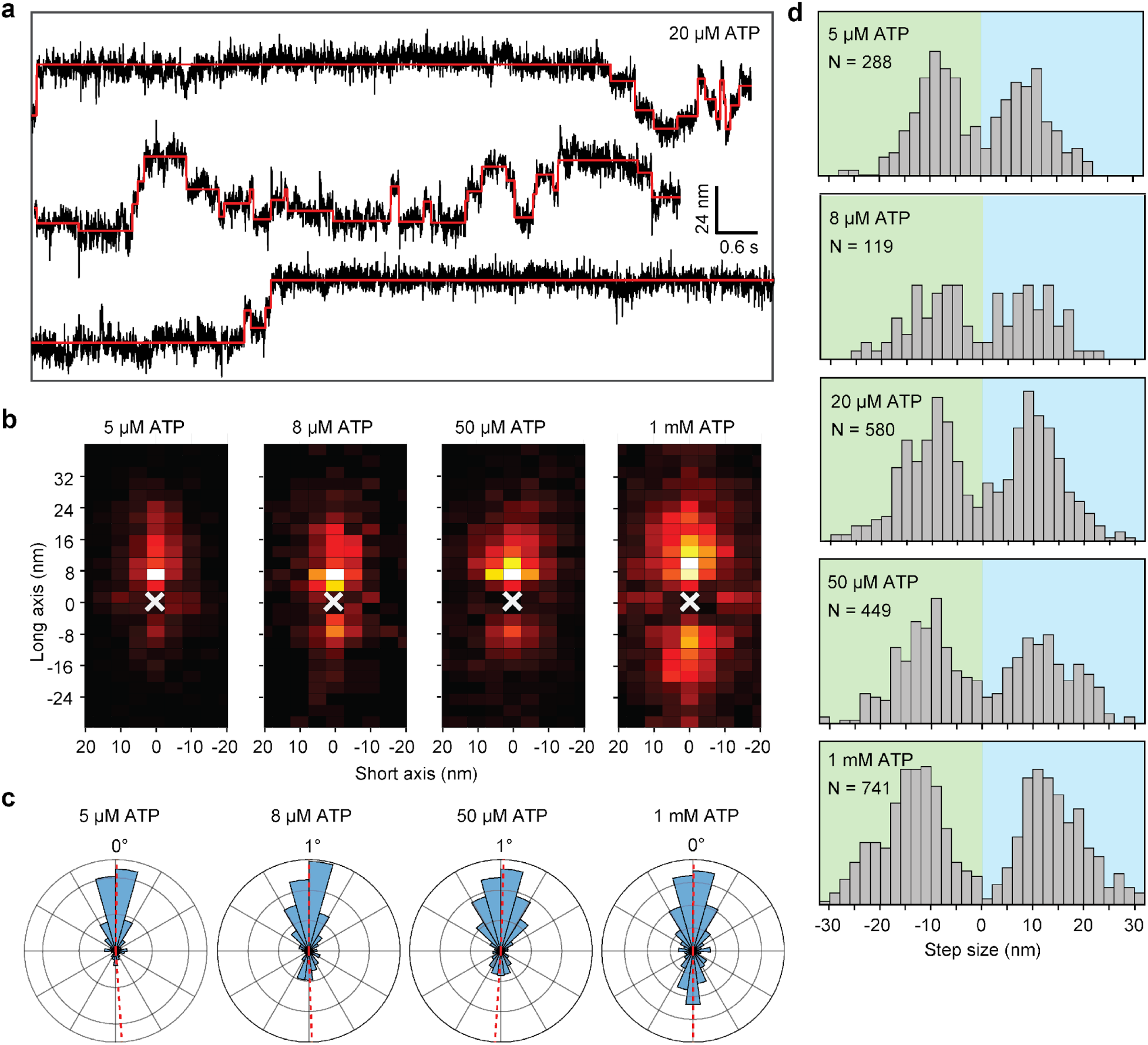
Two-dimensional step analysis of MINFLUX trajectories. **a,** Additional example trajectories of dynein stepping along the MT short-axis at 20 µM ATP. **b,** Heat maps show a high probability of dynein accessing neighboring protofilaments while stepping along the MT at different ATP conditions. **c,** Polar histograms of dynein’s stepping direction reveal the mean angular deviation of dynein from the MT minus-end (From left to right, n = 1020, 541, 1783, 2625 steps). **d,** The histogram of steps taken by dynein in the short-axis direction under different ATP concentrations.

**Extended Data Figure 6.**
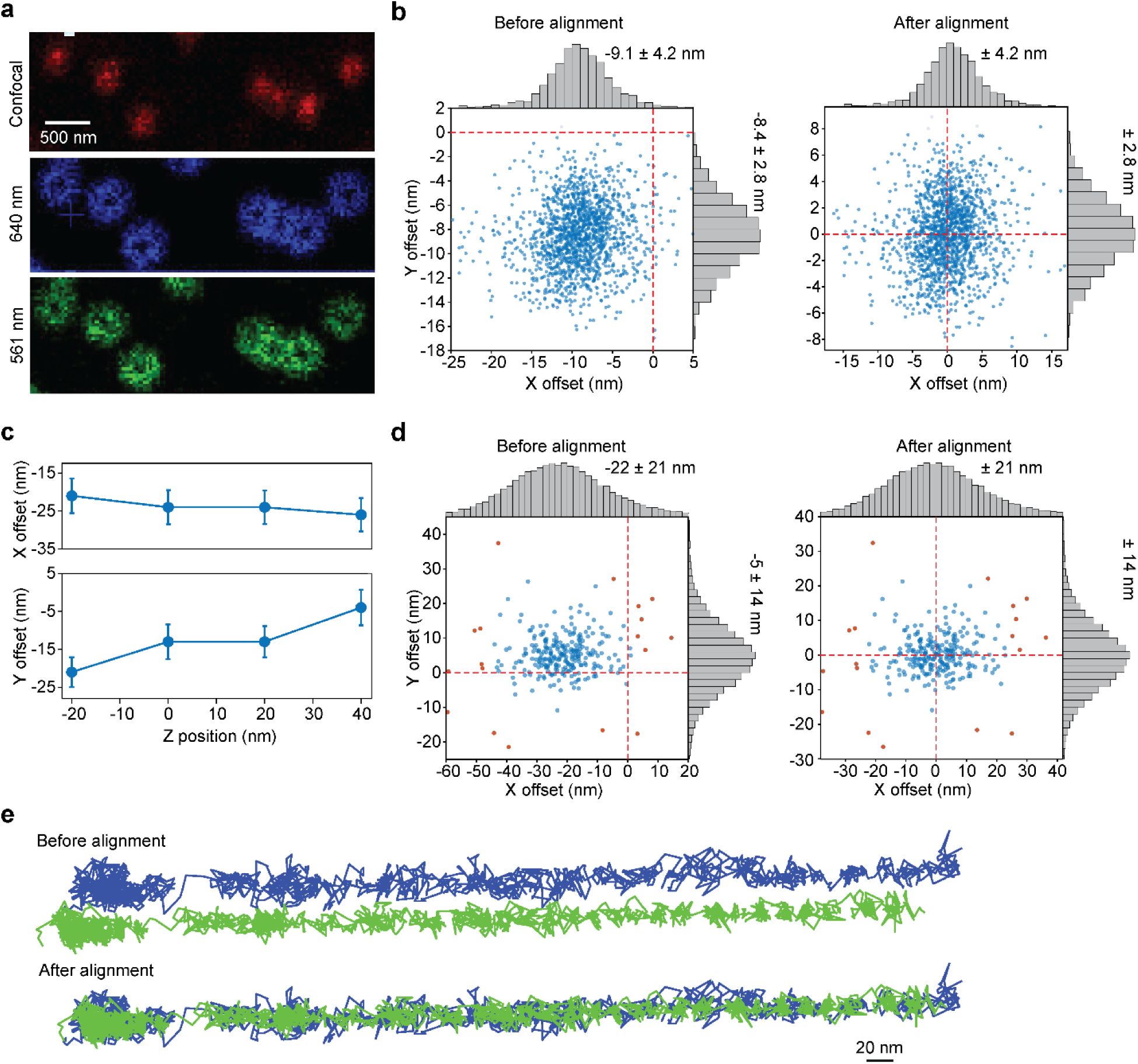
Registration of the two fluorescent channels in two-color MINFLUX. **a,** Tetraspec beads were illuminated by a toroidal confocal scanning beam, showing the shape of the local excitation minimum formed in both channels. **b,** Tetraspec bead localizations in both channels were used to calculate the offset in *x* and *y* directions (mean ± s.d.; n = 1606 localizations). The s.e.m. of the offset is 0.26 nm. The mean offset in both directions was corrected to align the two channels with ∼4 nm precision. **c,** The offset between the two channels varies substantially with the z-position of the objective (mean ± s.d.; from left to right, n = 323, 370, 335, 285 localizations). **d,** Two-color MINFLUX trajectories of dynein walking on a single MT were pooled to calculate the mean offset in *x* and *y* directions (n = 1606 localizations). The s.e.m. of the offset is 1.3 nm. **e,** An example two-color MINFLUX trajectory of kinesin-1 before and after alignment.

**Extended Data Figure 7.**
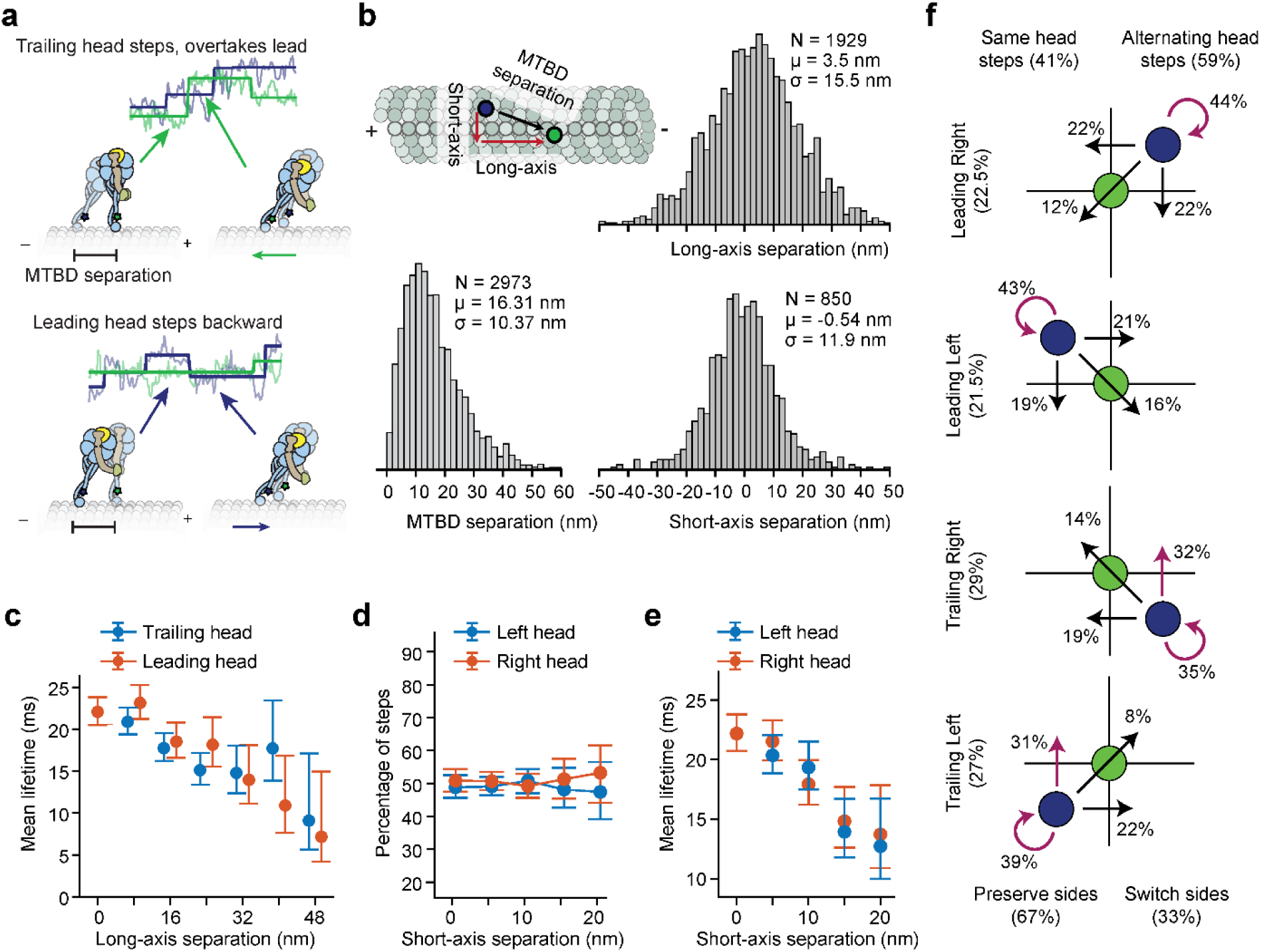
The analysis of two-color MINFLUX trajectories. **a,** Exemplary trajectories and schematics depicting the uncoordinated stepping behavior of dynein. **b,** The distribution of the distance on the *xy* plane between the two MTBDs of dynein during stepping. **c,** Mean lifetime of steps taken by either the leading or trailing head along the MT long axis under different MTBD separations (from left to right, n = 652, 433, 249, 106, 53 steps for the trailing and 661, 500, 297, 146, 64, 24 steps for the leading head). **d-e,** Percentage of steps (d) and mean lifetime (e) of steps taken by the right or left head under different MTBD separations in the MT short axis (from left to right, n = 633, 356, 134, 64 steps for the left, and 808, 619, 359, 128, 59 steps for the right head). **f,** A dissection of the relative position of the heads before and after a step. The arrows represent the relative position of the stepping head (blue) to its MTBD-bound partner (green) after a step. The trailing head is more likely to overtake the lead than the leading head to step backward to move to the trailing position.

**Extended Data Figure 8.**
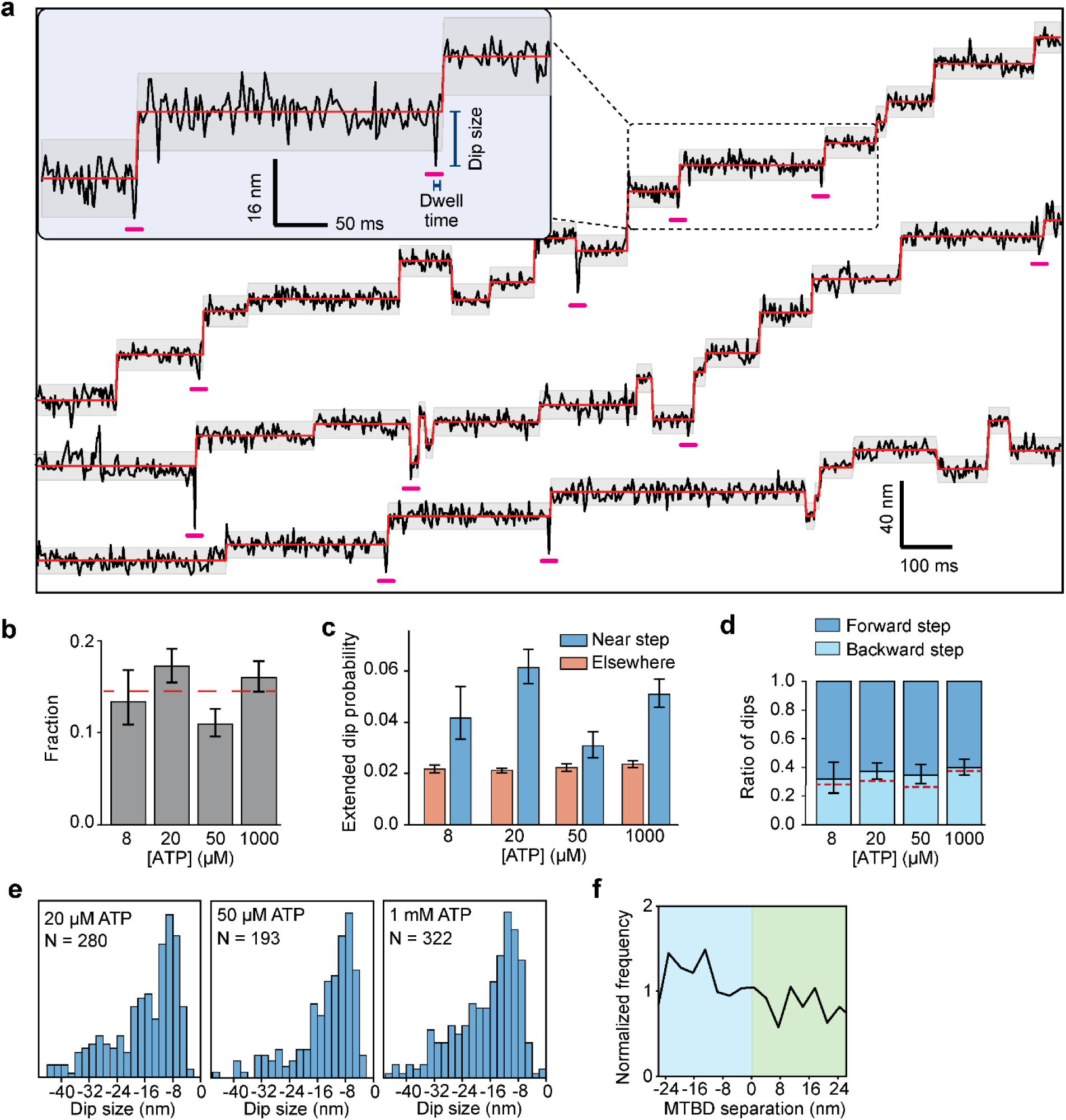
Analysis of dips in trajectories. **a,** Exemplary stepping trajectories on the MT long axis recorded at 2.5 ms temporal resolution at 20 µM ATP. Dips are underscored in pink. **b,** The probability of a step preceded by a dip is independent of ATP concentration. Error bars represent s.e.m. The red dashed line represents the average fraction of steps with dips across all tested conditions. **c,** The probability of a dip event occurring within 3 frames before a step or elsewhere during the dwell. Error bars represent ±95% c.i. **d,** The ratio of dips that occur before a forward or a backward step. The centerline and whiskers represent the mean and ±95% c.i.. The dashed red line represents the probability of backward steps at given ATP concentrations (from left to right, n = 485, 1614, 1865, 2326 steps). **e,** The histogram of the size of dips that occur within 3 frames before a step at different ATP concentrations. **f,** The normalized frequency of the dips in two-color MINFLUX traces as a function of interhead separation. Positive separation indicates steps taken by the leading head.

**Extended Data Figure 9.**
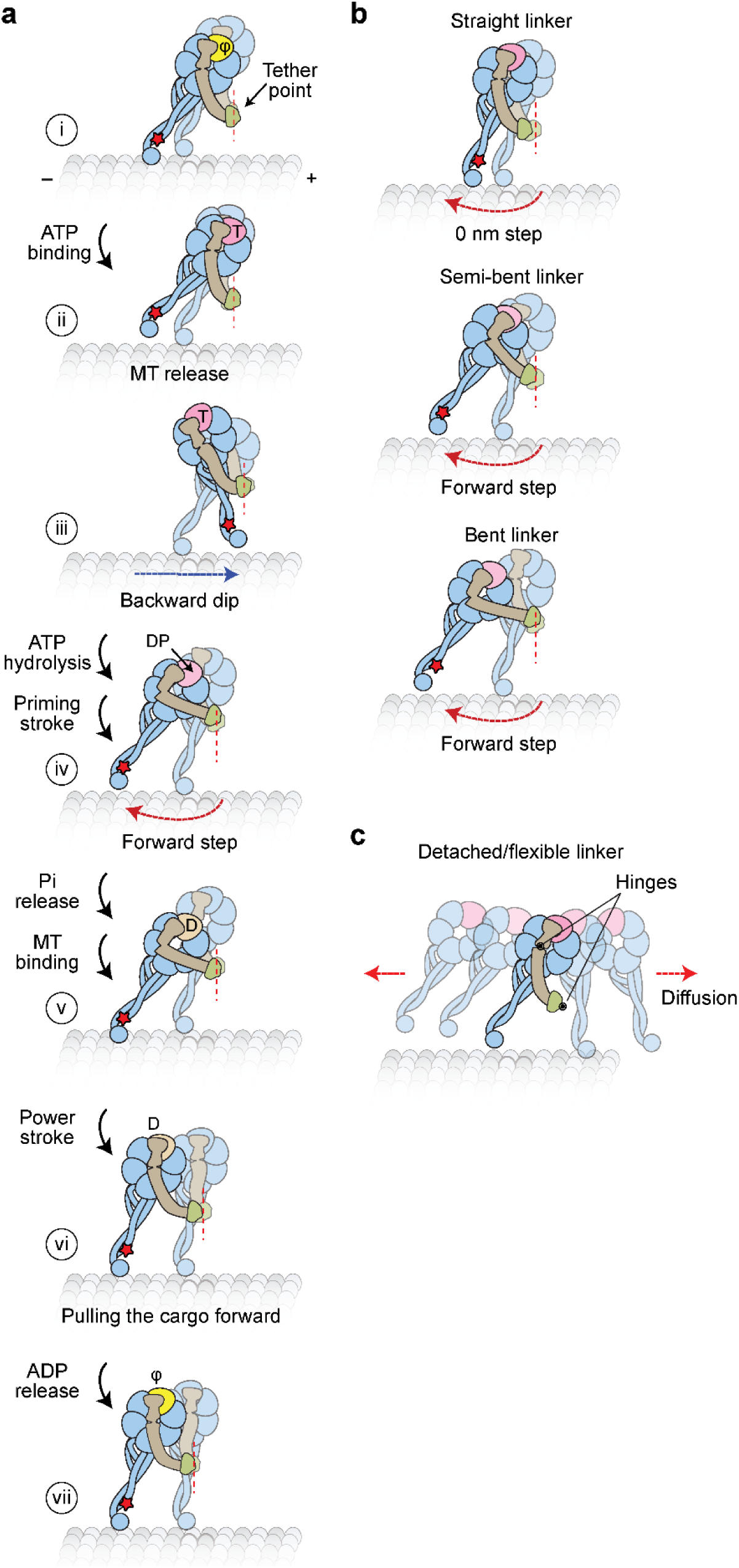
Model for the productive ATPase cycle and transient dips of dynein stepping. **a,** Updated model for the mechanochemical cycle of dynein. (i) When AAA1 is in the nucleotide-free state, a dynein head is bound to the MT, and the linker is in a straight conformation. (ii) ATP binding to AAA1 triggers the release of the motor from the MT. (iii) After MT release, the dynein head moves toward the tethering point near the dimerization domain and undergoes tethered diffusion, resulting in backward dips observed in dynein trajectories. (iv) After ATP hydrolysis, the priming stroke of the linker pushes the MTBD towards the MT minus end. (v) Dynein rebinds the MT and releases inorganic phosphate (Pi). (vi) The Pi release stabilizes the straight conformation of the linker, exiting the AAA ring near AAA4, and dynein pulls its cargo in the forward direction. (vii) Upon ADP release, the linker further clockwise, exiting the AAA ring near AAA5, and the motor resets for the next ATPase cycle. **b,** Different conformations of the linker in the ADP.Pi state may contribute to the high variability of the dynein step size. The linker can orient in straight, semi-bent, and bent conformations or detach from the AAA+ ring. Straightening of the linker before MT rebinding may result in no net displacement and account for 0 nm steps. Semi-bent and bent conformations push the MTBD towards the minus-end, which may be responsible for the small minus-end-directed bias in the dynein step size. **c,** Detachment of the linker from the AAA+ ring increases the degrees of freedom and results in a larger diffusional search, which may account for the high variability of the dynein step size distribution.

**Extended Data Figure 10.**
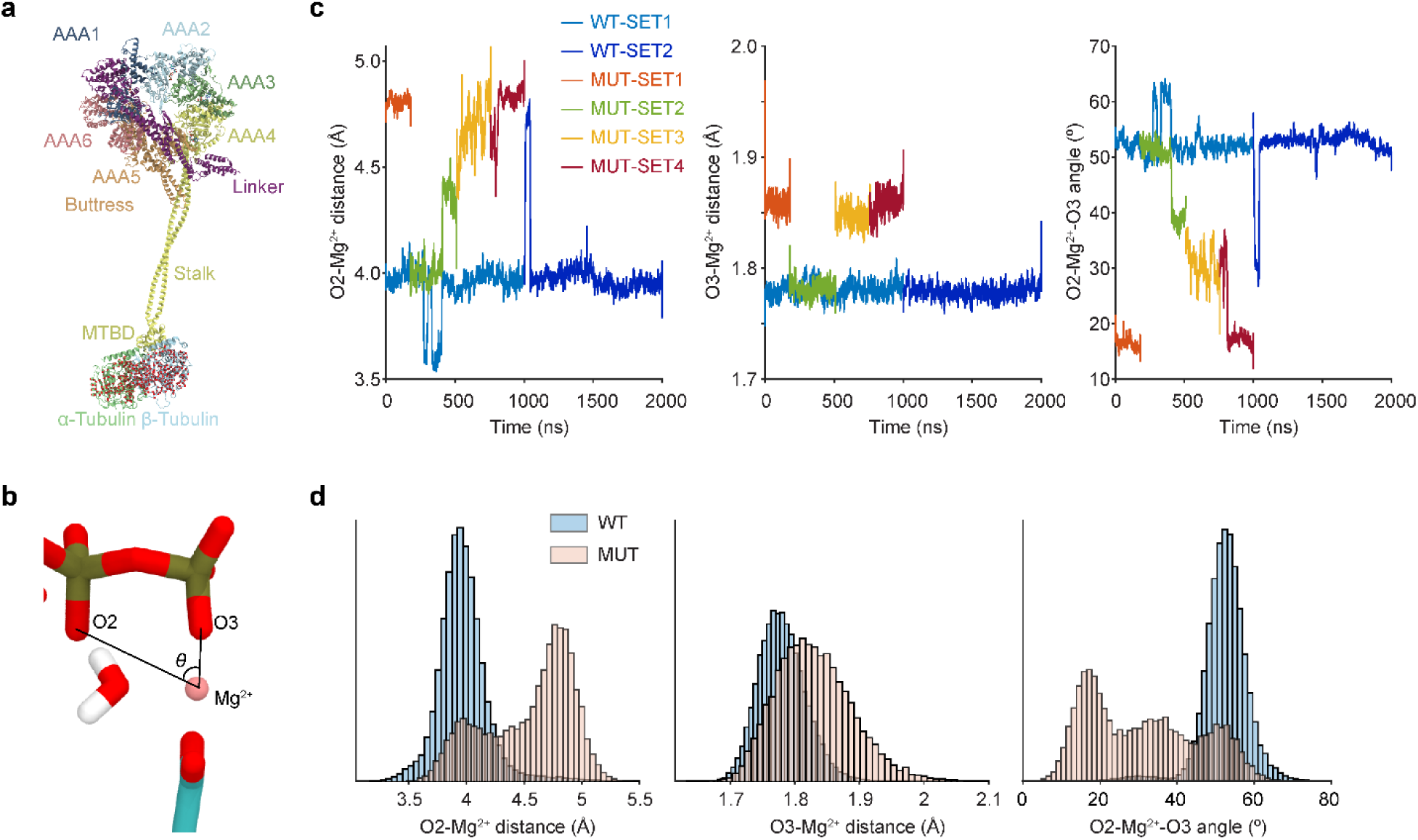
MD simulations show the D1848E mutation disrupts the coordination of ATP by Mg²⁺ at AAA1. **a,** The structural model of dynein bound to tubulin in MD simulations (see Methods). Red dots in tubulin represent constraints introduced to Cα atoms to maintain the tubulin conformation on MTs. **b,** A cartoon depiction shows the distances and the angle between O2, Mg²⁺, and O3. **c,** Time evolution of Mg²⁺ coordination: O2–Mg²⁺ distances (left), O3–Mg²⁺ distances (middle), and O2–Mg²⁺–O3 angle (right) are shown for WT and D1848E simulations. WT trajectories maintain a stable coordination geometry, whereas the mutant shows substantial fluctuations and loss of defined geometry. **d,** Normalized distributions of O2–Mg²⁺ distances (left), O3–Mg²⁺ distances (middle), and O2–Mg²⁺–O3 angles (right) highlight narrow, well-defined peaks in the WT compared with the broader, destabilized distributions in D1848E. In the WT system, the O2–Mg²⁺ distance remained consistently 3.6–3.8 Å, while the O3–Mg²⁺ distance and O2–Mg²⁺–O3 angle were tightly distributed around 1.8 Å and 40–50°, respectively. In contrast, the D1958E mutant exhibited a marked displacement of Mg²⁺, with O2–Mg distances fluctuating beyond 4.5 Å and broader, destabilized distributions for both the O3–Mg distance and O2–Mg–O3 angle. The displacement of Mg²⁺ positions O3 between Mg²⁺ and O2, effectively abolishing the O2–Mg²⁺ coordination and reducing the efficiency of ATP hydrolysis.

**Extended Data Figure 11.**
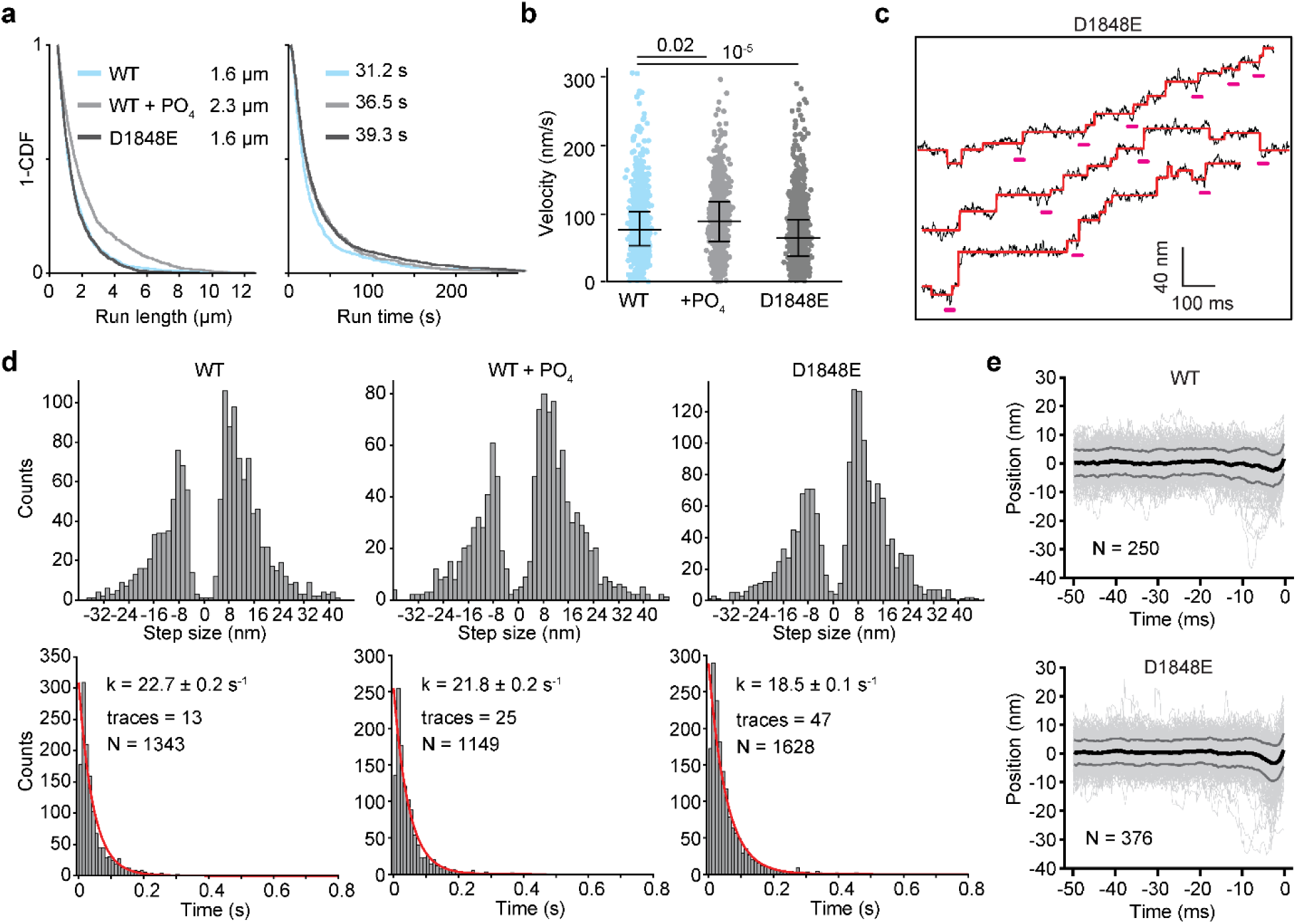
Motility and stepping characteristics of WT and mutant dynein. **a,** The run length and run time of WT, WT with 30 mM PO_4_, and D1848E mutant dynein in 1 mM ATP (n = 955, 861, 1201 motors, respectively). **b,** Distribution of velocities of WT dynein, WT with 30 mM PO_4_ (+PO_4_), and D1848E at 1 mM ATP in one of the four biological replicates in Fig. 5b. The centerline and whiskers represent mean and s.d., respectively (from left to right, n = 956, 865, 1202 motors). *P* values were calculated from a two-tailed t-test. **c,** Additional example trajectories of D1848E at 2.9 nm spatial and 0.45 ms temporal resolution at 1 mM ATP. Red lines represent a fit to a step finding algorithm. Pink underlines represent detected dips. **d,** Step size (top) and dwell time (bottom) histograms of WT, WT with 30 mM PO_4_, and D1848E stepping at 1 mM ATP. Dashed red curves represent a single exponential fit to calculate the stepping rate (±s.e.). **e,** The cumulative overlay of the dwell positions WT and D1848E before a step (t = 0 s) demonstrates the presence of a backward dip. Black and grey curves represent mean ± 2σ, respectively. The FWHM of the dips in D1848E was 3.85 ms, similar to the dip lifetime of WT dynein (Figs. 4b and d).

**Extended Data Table 1.**
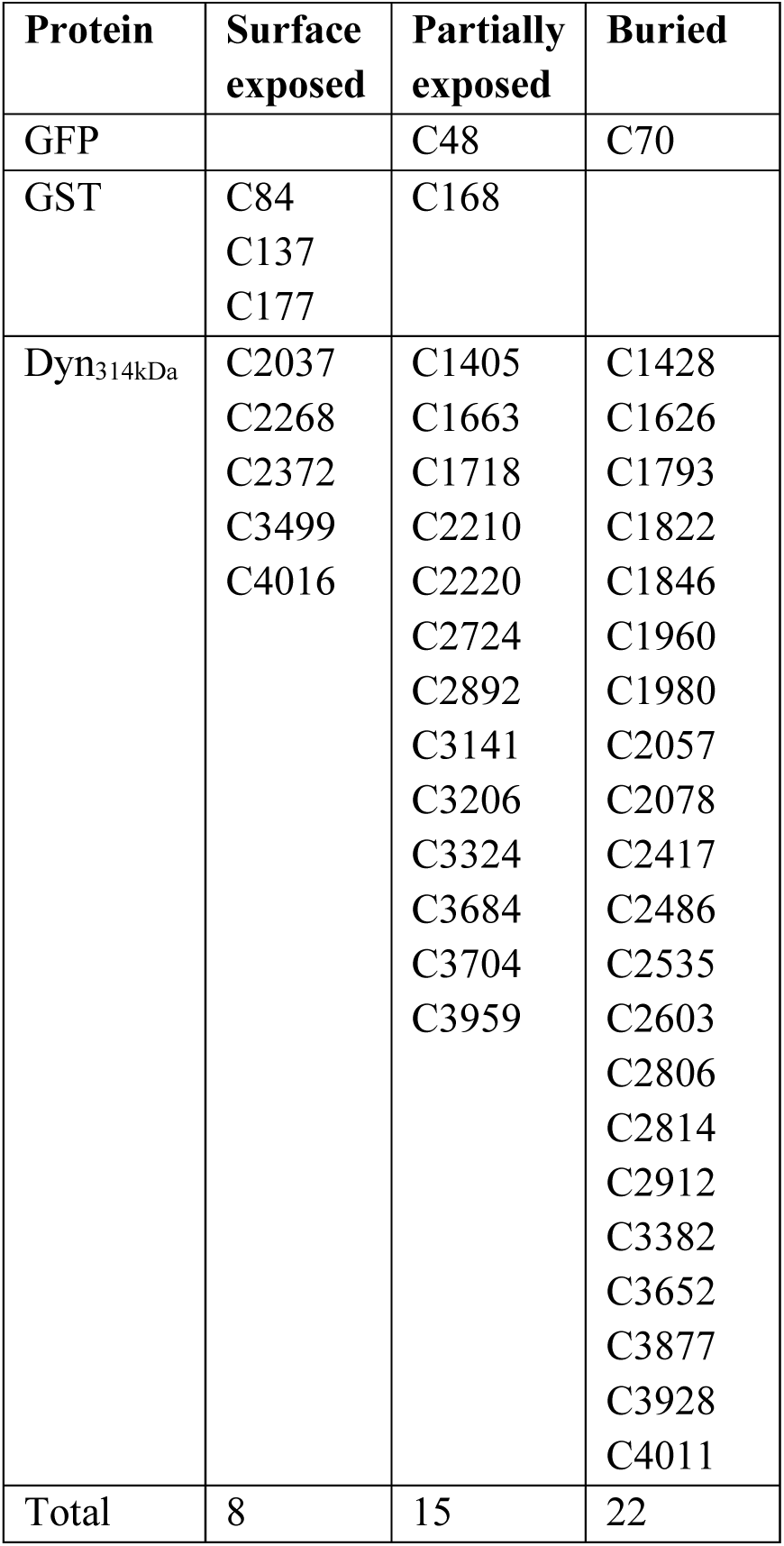
Characterization of cysteines in truncated dynein. Cysteines of the dynein motor domain were categorized into three groups based on the available X-ray structure of yeast dynein (PDB ID: 4AKG)^27^. The surface-exposed cysteines were accessible to surrounding water over 5 Å, partially exposed cysteines were solvent accessible to over 2.5 Å, and buried cysteines to less than 2.5 Å. Surface-exposed cysteines of the GST-dimerization were also removed. SpyCatcher and SpyTag do not contain any cysteines and are therefore omitted from this table.

**Extended Data Table 2.** Testing of unique cysteine for labeling Dyn_CLM_ at its MTBD. Protein expression and motility properties of Dyn_CLM_ constructs mutated with a single surface-exposed cysteine at the MTBD, compared to WT Dyn_314kDa_. The mutations were introduced to the GST-Dyn_CLM_ construct. Assays were performed in 1 mM ATP. In velocity measurements, N = 50 motors for each construct (two technical replicates). The most processive and well-expressing motor (Q3231C) was used in this study. The denaturing gel used for quantifying protein expression levels is shown in Extended Data Fig. 1.

| **Construct Name** | **Expression Level** | **Velocity (nm/s) (mean ± s.d.)** | **Motility** |
| --- | --- | --- | --- |
| WT | High | $80 \pm 23$ | High |
| S3097C | Low | $100 \pm 19$ | Low |
| E3107C | Low | $58 \pm 14$ | Low |
| Q3147C | Low | $10 \pm 5$ | High |
| H3169C | High | $80 \pm 22$ | Medium |
| H3175C | Low | $83 \pm 22$ | Medium |
| Q3179C | High | $70 \pm 13$ | Medium |
| K3182C | Medium | $73.3 \pm 17$ | Low |
| S3190C | Medium | $78.6 \pm 24$ | Low |
| Q3211C | Medium | $76.6 \pm 16$ | Low |
| **Q3231C** | **Medium** | **$80 \pm 19$** | **High** |
| K3234C | None | - | - |
| K3242C | Medium | $50 \pm 12$ | Medium |

**Extended Data Table 3.** Yeast strains and plasmids used in this study. UPF1 was knocked out of the CY1 genome (MATa; his3-11,15; ura3-1; leu2-3,112; ade2-1; trp1-1) using the KanMX6 marker. CY1 was used as an acceptor strain for exogenous expression of cys-lite dynein constructs in a pYES plasmid backbone. HALO (DHA) is a genetic tag used for labeling the protein with a fluorescent dye.

| Strain Name | Description | Genotype | Database | Reference |
| --- | --- | --- | --- | --- |
| WT Dynein | GFP-GST-Dyn1<sub>314kDa</sub> | *MATa; his3-11,15; ura3-1; leu2-3,112; ade2-1; trp1-1; PEP4::HIS5; PRB1D pDyn-pGAL-ZZ-TEV-GFP-3XHA-GST-DYN1<sub>314kDa</sub>-gsDHA:Kan* | CY31 | <sup>23</sup> |
| Tail-labeled Dynein | DHA-GST-Dyn1<sub>314kDa</sub> | *MATa; his3-11,15; ura3-1; leu2-3,112; ade2-1; trp1-1; PEP4::HIS5; PRB1D pDyn-pGAL-ZZ-TEV-gsDHA-GST-DYN1<sub>314kDa</sub>* | VY268 | <sup>23</sup> |

**Extended Data Table 3 | Yeast strains and plasmids used in this study.**
| Plasmid Name | Description | Database | Reference |
| --- | --- | --- | --- |
| GST<sub>CLM</sub>-Dyn | *pDyn-Ura3-pGAL-ZZ-TEV-GFP-3XHA-GST<sub>C84S,C137S,C177S</sub>-D6-Dyn1<sub>314kDa</sub>* | CY392 | This study |
| GST<sub>CLM</sub> - Dyn<sub>CLM</sub> | *pDyn-Ura3-pGAL-ZZ-TEV-GFP-3XHA-GST<sub>C84S,C137S,C177S</sub>-Dyn1<sub>314kDa</sub>, C2037S, C2268S, C2372S, C3499S, C4016S* | CY393 | This study |
| GST<sub>CLM</sub>-Dyn<sub>CLM</sub> Q3231C | *pDyn-Ura3-pGAL-ZZ-TEV-GFP-3XHA-GST<sub>C84S,C137S,C177S</sub> - Dyn1<sub>314kDa</sub>, C2037S, C2268S, C2372S, C3499S, C4016S, Q3231C* | CY486 | This study |
| GST<sub>CLM</sub>-Dyn<sub>CLM</sub> Q3231C, D1848E | *pDyn-Ura3-pGAL-ZZ-TEV-GFP-3XHA-GST<sub>C84S,C137S,C177S</sub> - Dyn1<sub>314kDa</sub>, C2037S, C2268S, C2372S, C3499S, C4016S, Q3231C, D1848E* | JS36 | This study |
| SpyCatcher-Dyn<sub>CLM</sub> | *pDyn-Ura3-pGAL-ZZ-TEV-GFP-3XHA- SpyCatcher-Dyn1<sub>314kDa</sub>, C2037S, C2268S, C2372S, C3499S, C4016S* | CY483 | This study |
| SpyTag-Dyn<sub>CLM</sub> | *pDyn-Ura3-pGAL-ZZ-TEV-GFP-3XHA- SpyTag-Dyn1<sub>314kDa</sub>, C2037S, C2268S, C2372S, C3499S, C4016S* | CY487 | This study |
| SpyTag-Dyn<sub>CLM</sub> Q3231C | *pDyn-Ura3-pGAL-ZZ-TEV-GFP-3XHA- SpyTag-Dyn1<sub>314kDa</sub>, C2037S, C2268S, C2372S, C3499S, C4016S, Q3231C* | CY488 | This study |
| SpyCatcher-Dyn<sub>CLM</sub> Q3231C | *pDyn-Ura3-pGAL-ZZ-TEV-GFP-3XHA- SpyCatcher-Dyn1<sub>314kDa</sub>, C2037S, C2268S, C2372S, C3499S, C4016S, Q3231C* | CY489 | This study |

**Extended Data Table 4.**
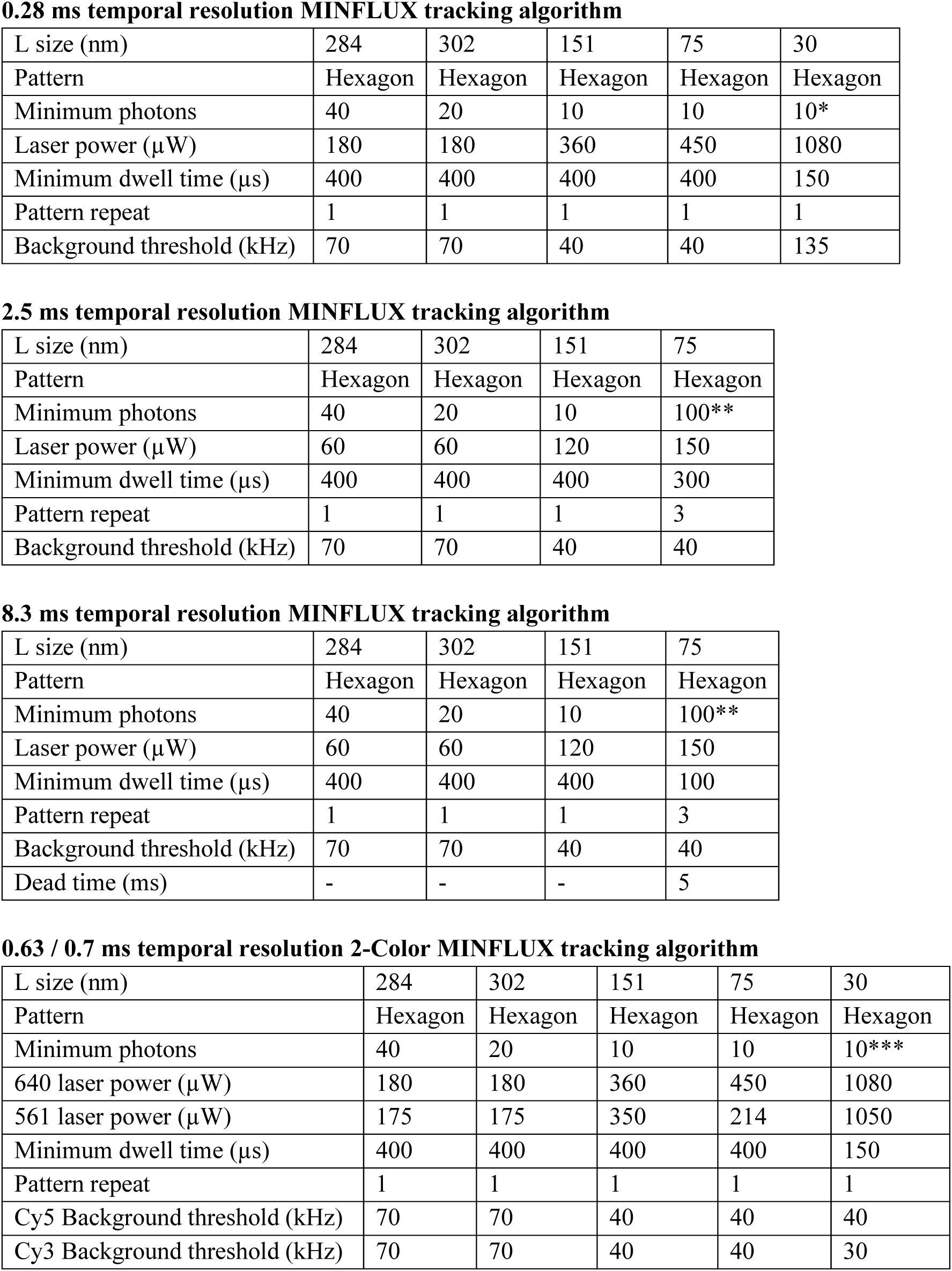
List of MINFLUX parameters used to track dynein at different temporal resolutions. 10* and 100** represent the minimum photon counts needed for the next iteration. Laser power was optimized to obtain an average number of 30*, 140**, and 15*** photons, respectively. The 0.28 ms temporal resolution and two-color algorithm also required increasing the pinhole to 0.97 Airy units. All reported powers are measured at the periscope and coupled to the fiber at 35% efficiency.

**0.28 ms temporal resolution MINFLUX tracking algorithm**
|  |  |  |  |  |  |
| --- | --- | --- | --- | --- | --- |
| L size (nm) | 284 | 302 | 151 | 75 | 30 |
| Pattern | Hexagon | Hexagon | Hexagon | Hexagon | Hexagon |
| Minimum photons | 40 | 20 | 10 | 10 | 10* |
| Laser power (μW) | 180 | 180 | 360 | 450 | 1080 |
| Minimum dwell time (μs) | 400 | 400 | 400 | 400 | 150 |
| Pattern repeat | 1 | 1 | 1 | 1 | 1 |
| Background threshold (kHz) | 70 | 70 | 40 | 40 | 135 |

**2.5 ms temporal resolution MINFLUX tracking algorithm**
|  |  |  |  |  |
| --- | --- | --- | --- | --- |
| L size (nm) | 284 | 302 | 151 | 75 |
| Pattern | Hexagon | Hexagon | Hexagon | Hexagon |
| Minimum photons | 40 | 20 | 10 | 100** |
| Laser power (μW) | 60 | 60 | 120 | 150 |
| Minimum dwell time (μs) | 400 | 400 | 400 | 300 |
| Pattern repeat | 1 | 1 | 1 | 3 |
| Background threshold (kHz) | 70 | 70 | 40 | 40 |

**8.3 ms temporal resolution MINFLUX tracking algorithm**
|  |  |  |  |  |
| --- | --- | --- | --- | --- |
| L size (nm) | 284 | 302 | 151 | 75 |
| Pattern | Hexagon | Hexagon | Hexagon | Hexagon |
| Minimum photons | 40 | 20 | 10 | 100** |
| Laser power (μW) | 60 | 60 | 120 | 150 |
| Minimum dwell time (μs) | 400 | 400 | 400 | 100 |
| Pattern repeat | 1 | 1 | 1 | 3 |
| Background threshold (kHz) | 70 | 70 | 40 | 40 |
| Dead time (ms) | - | - | - | 5 |

**Extended Data Table 4 | List of MINFLUX parameters used to track dynein at different temporal resolutions.**
|  |  |  |  |  |  |
| --- | --- | --- | --- | --- | --- |
| L size (nm) | 284 | 302 | 151 | 75 | 30 |
| Pattern | Hexagon | Hexagon | Hexagon | Hexagon | Hexagon |
| Minimum photons | 40 | 20 | 10 | 10 | 10*** |
| 640 laser power (μW) | 180 | 180 | 360 | 450 | 1080 |
| 561 laser power (μW) | 175 | 175 | 350 | 214 | 1050 |
| Minimum dwell time (μs) | 400 | 400 | 400 | 400 | 150 |
| Pattern repeat | 1 | 1 | 1 | 1 | 1 |
| Cy5 Background threshold (kHz) | 70 | 70 | 40 | 40 | 40 |
| Cy3 Background threshold (kHz) | 70 | 70 | 40 | 40 | 30 |

## Notes

### Competing Interest Statement

The authors have declared no competing interest.

### Summary of Updates

We performed two-color MINFLUX tracking of dynein and revealed how the stepping of the two heads is partially coordinated through intramolecular tension as the heads separate from each other. We designed an ATP hydrolysis mutant of the principal ATPase site (AAA1) of dynein to understand the backward dips we observed in the dynein mechanochemical cycle. The new experiments and modeling studies revealed which step of the mechanochemical cycle (ATP hydrolysis) produces a net step. This characterizes these dips as a key intermediate of the step of dynein which resolves a major controversy in the field.

